# F-actin-rich territories coordinate apoptosome assembly and caspase activation during intrinsic apoptosis

**DOI:** 10.1101/2022.08.05.502994

**Authors:** Virginia L King, Kenneth G Campellone

## Abstract

The actin cytoskeleton is a ubiquitous participant in cellular functions that maintain viability, but how it controls programmed cell death processes is not well understood. Here we show that in response to DNA damage, human cells form juxtanuclear F-actin-rich territories that coordinate the organized progression of apoptosome assembly to caspase activation. These cytoskeletal compartments are created by the actin nucleation factors JMY, WHAMM, and the Arp2/3 complex, and they exclude proteins that inhibit JMY and WHAMM activity. Within the territories, JMY localization overlaps with punctate structures containing the core apoptosome components cytochrome *c* and Apaf-1. The F-actin-rich areas also encompass initiator caspase-9 and clusters of a cleaved form of executioner caspase-3, while restricting accessibility of the caspase inhibitor XIAP. The clustering and potency of caspase-3 activation are positively regulated by the amount of actin polymerized by JMY and WHAMM. These results indicate that JMY-mediated actin reorganization functions in apoptotic signaling by coupling the biogenesis of apoptosomes to the localized processing of caspases.

## INTRODUCTION

Actin, one of the most ubiquitous proteins in eukaryotic cells, is polymerized into filaments that can generate forces, create scaffolds, and act as tracks for motor proteins, making the actin cytoskeleton a dynamic and essential participant in numerous cellular functions. The formation, organization, and turnover of filamentous (F-) actin is regulated by a multitude of actin-binding proteins (Pollard, 2016). To create branched actin networks, the heteroheptameric Arp2/3 complex nucleates actin filaments in collaboration with nucleation-promoting factors from the Wiskott-Aldrich Syndrome Protein (WASP) family (Campellone and Welch, 2010). The mammalian WASP family is composed of 9 members that stimulate Arp2/3-mediated actin assembly at distinct cellular localizations in response to different signals (Rotty et al., 2013; Alekhina et al., 2017; Kabrawala et al., 2020). Functions for the actin cytoskeleton during cell motility, adhesion, and division have been extensively characterized (Swaney and Li, 2016; Svitkina, 2018; Pollard and O’Shaughnessy, 2019), and newer activities for nucleation factors in membrane trafficking and chromatin dynamics are emerging (Caridi et al., 2019; Chakrabarti et al., 2021), but their roles in programmed cell death remain understudied.

The normal progression of many organismal processes, such as embryogenesis, tissue turnover, and tumor suppression, relies on the tightly regulated form of cell death called apoptosis (Kerr et al., 1972; Galluzzi et al., 2018). Apoptosis can be driven by intrinsic mitochondria-mediated and extrinsic receptor-mediated death mechanisms that converge on a terminal execution program (von Karstedt et al., 2017; Singh et al., 2019). Intrinsic pathways are initiated in response to intracellular damage and are characterized by mitochondrial outer membrane permeabilization resulting in the export of apoptogenic proteins, including cytochrome *c* (cyto *c*), from the mitochondria to the cytosol (Tait and Green, 2013; Bock and Tait, 2020). Once cytosolic, cyto *c* can interact with the Apoptotic protease activating factor-1 (Apaf-1) to instigate the assembly of macromolecular platforms called apoptosomes (Riedl and Salvesen, 2007). Apoptosomes mediate the multimerization and proteolytic processing of initiator and executioner caspases (Bratton and Salvesen, 2010; Yuan and Akey, 2013; Dorstyn et al., 2018), the latter of which eventually cleave multiple proteins in the cytosol and nucleus to drive a cell suicide program (Green and Llambi, 2015; Julien and Wells, 2017). While the structural and biophysical properties of apoptosomes have come into focus *in vitro* (Rodriquez and Lazebnik, 1999; Zou et al., 1999; Yin et al., 2006; Yuan et al., 2011; Yuan et al., 2013; Hu et al., 2014; Zhou et al., 2015; Cheng et al., 2016; Li et al., 2017), their biological characteristics within cells remain less clear.

Cytoskeletal studies of apoptotic cell death were initially linked to the late-stage changes in cell adherence and morphology that are controlled by actin filament disassembly, rearrangement, and even cleavage by caspases (Gourlay and Ayscough, 2005; Desouza et al., 2012). Actin is also recruited to mitochondria earlier in apoptosis at around the time of mitochondrial permeabilization (Tang et al., 2006; Wang et al., 2008; Rehklau et al., 2012), and the actin turnover machinery has been implicated in controlling intrinsic apoptosis at multiple steps. The mitochondrial localization of cofilin, which depolymerizes actin filaments, influences the release of cyto *c* (Chua et al., 2003; Zhu et al., 2006; Klamt et al., 2009). Moreover, the F-actin-severing protein gelsolin may regulate mitochondrial membrane integrity, be cleaved by caspases, stimulate depolymerization, and allow chromatin fragmentation (Kothakota et al., 1997; Kusano et al., 2000; Ahn et al., 2003; Chhabra et al., 2005).

More recently, roles for actin assembly proteins as active participants in DNA damage-induced apoptosis have been uncovered. One of the WASP-family members, JMY, was discovered as a cofactor that affects the function of the tumor suppressor protein and transcription factor p53 (Shikama et al., 1999). Transient overexpression and depletion experiments suggest that JMY can promote cell death by enhancing p53-mediated transcription of pro-apoptotic genes (Shikama et al., 1999; Coutts et al., 2009). However, the activity of JMY in transcriptional regulation via p53 may not be the primary apoptotic driver following DNA damage, as the genetic deletion of JMY does not impact nuclear p53 modification or p53-driven transcription of genes that canonically lead to a proliferation arrest or mitochondrial permeabilization (King et al., 2021).

Under normal cellular growth conditions, the abundance of both JMY and p53 is modulated by the ubiquitin ligase Mdm2, which promotes their proteasomal degradation (Coutts et al., 2007). JMY shuttles between the nucleus and cytosol (Coutts et al., 2009; Zuchero et al., 2012), and although it may have pro-apoptotic functions in the nucleus, its most extensively characterized activities are as a cytosolic actin nucleation factor (Zuchero et al., 2009). In fact, following DNA damage, JMY, its closest homolog WHAMM, and the Arp2/3 complex are necessary for rapid pro-apoptotic responses in the cytoplasm (King et al., 2021). Cells lacking JMY or WHAMM exhibit kinetic delays or deficiencies in the mitochondrial export of cyto *c*, activation of caspases, and ultimately the terminal cleavage events throughout the cell (King et al., 2021). In further support of the importance of actin assembly in intrinsic death pathways, apoptotic cells form cytoplasmic structures that contain JMY, F-actin, cyto *c*, and active executioner caspase-3 (King et al., 2021).

Collectively, these studies indicate that the actin cytoskeleton is an important player in the transition from mitochondrial permeabilization to executioner caspase activation. However, the underlying mechanisms by which it impacts progression through this crucial period of intrinsic apoptotic signaling are unclear. In the current study, we characterized the intracellular organization and function of multiple elements of the actin regulatory and apoptotic machinery and discovered that JMY-mediated actin polymerization creates a cytosolic F-actin-rich territory that compartmentalizes the apoptosome biogenesis process and promotes an efficient localized activation of caspases.

## RESULTS

### JMY reorganization into a cytosolic F-actin-rich territory is a more prominent step of the apoptotic response than its import into the nucleus

Our recent study showed that JMY, cyto *c*, active executioner caspases, and actin filaments form clusters of cytosolic puncta following DNA damage in multiple cell lines (King et al., 2021). To evaluate the timing of JMY puncta formation, we treated U2OS osteosarcoma cells, which are commonly employed in studies of apoptosis, with etoposide, a topoisomerase II inhibitor that induces dsDNA breaks. We then visualized the localization of endogenous JMY or stably-expressed GFP-tagged JMY (Figure S1) at regular intervals up to an 8h endpoint. As observed previously, both endogenous JMY and GFP-JMY formed intense juxtanuclear clusters of puncta in response to etoposide exposure (Figure 1A). At steady state ∼3% of cells contained such structures, and the proportion of JMY puncta-positive cells increased to 13% by 4h of etoposide treatment and reached ∼40% by 8h (Figure 1B), with each positive cell harboring a single zone of clustered JMY puncta. Staining with fluorescent phalloidin demonstrated that F-actin comprised the areas in-between and surrounding the JMY or GFP-JMY puncta to create a round overall structure with a typical diameter of 8-16µm (Figure 1A). These results extend earlier findings and show that the accumulation of JMY in punctate juxtanuclear structures increases during prolonged exposure to a DNA-damaging agent. We will refer to these cytoskeleton-associated clusters of JMY puncta as F-actin-rich territories.

**Figure 1.**
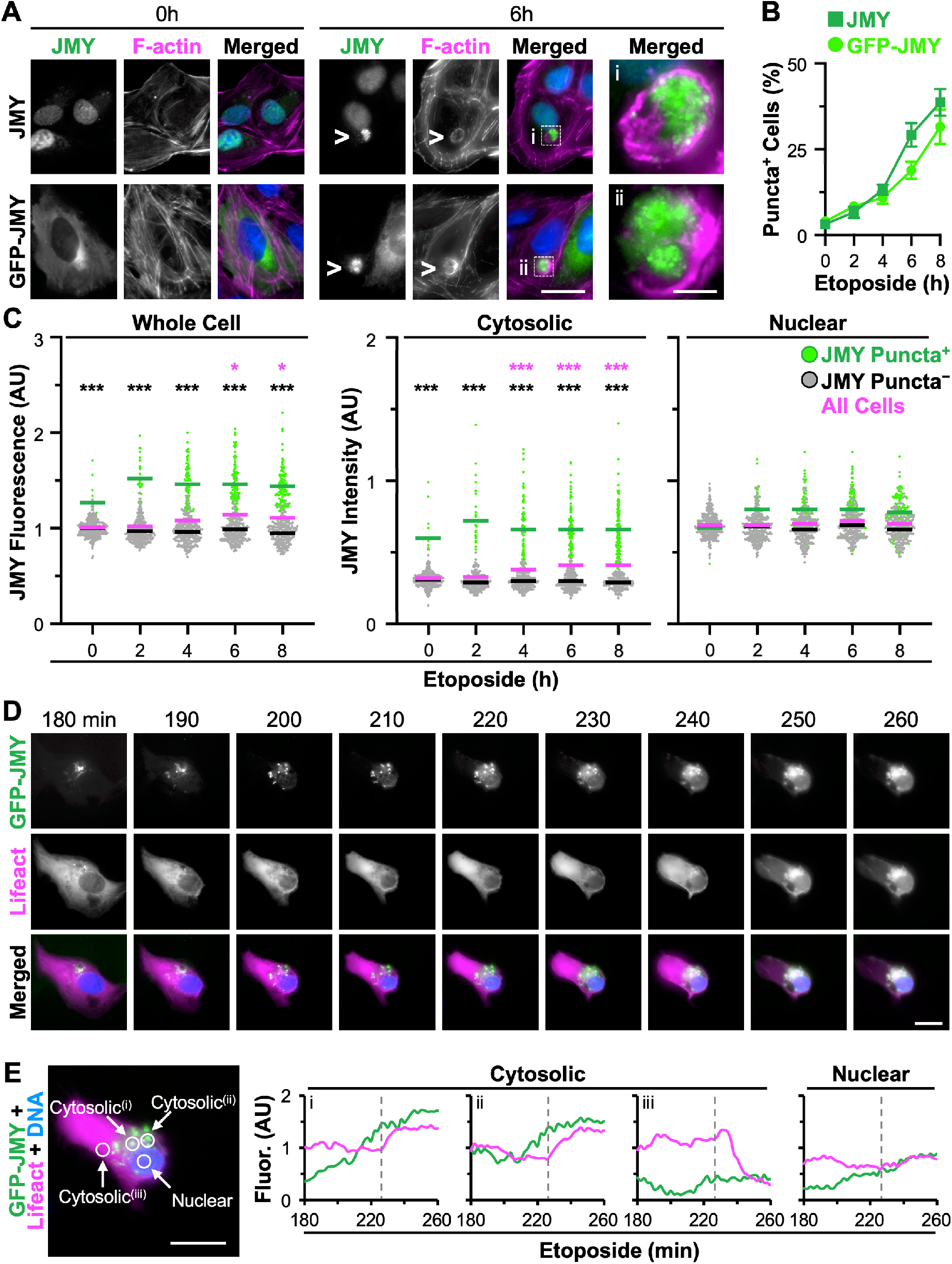
JMY forms cytosolic puncta prior to the assembly of juxtanuclear F-actin-rich territories. **(A)** U2OS cells were treated with 10μM etoposide for 0-8h before being fixed and stained with a JMY antibody (green) to visualize endogenous JMY, phalloidin to visualize F*-* actin (magenta), and DAPI to detect DNA (blue). U2OS cells expressing GFP-JMY (green) were also stained. Arrowheads and magnifications (i,ii) highlight examples of cytosolic JMY puncta. Scale bars, 25μm, 5μm. **(B)** The % of cells with JMY or GFP-JMY puncta was calculated. Each point represents the mean±SD from 3 experiments (n = 2,084-3,044 cells per point). **(C)** Whole cell, cytosolic, and nuclear JMY fluorescence values were measured in ImageJ. Each point represents 1 cell (n = 310-348 per timepoint) where green points are JMY puncta-positive cells and gray points are JMY puncta-negative cells. Green, black, and magenta bars are the mean values for the positive, negative, and total populations respectively, each from 3 experiments. Black significance stars refer to comparisons of the puncta-positive to puncta-negative population. Magenta stars refer to comparisons of the total population at each timepoint to the 0min timepoint. AU = arbitrary units. **(D)** U2OS cells expressing GFP-JMY (green) and Lifeact-mCherry (magenta) were treated with etoposide for 3h before incubation with Hoescht to visualize DNA (blue). Selected frames (taken from Video S1) show the formation of punctate GFP-JMY structures and an F-actin-rich territory. Scale bars, 20μm. **(E)** 5μm diameter circles were drawn within a cytosolic region containing punctate GFP-JMY (i,ii), a cytosolic region without GFP-JMY puncta (iii), or the nucleus, and the pixel intensity profiles for GFP-JMY and Lifeact-mCherry were measured. Dashed gray lines indicate the time when cytosolic F-actin territory formation begins. *p<0.05; ***p<0.001 (ANOVA, Tukey post-hoc tests).

It has been reported that JMY exhibits a predominantly cytoplasmic steady state localization in several cell lines (Coutts et al., 2009; Firat-Karalar et al., 2011; Schlüter et al., 2014; Hu and Mullins, 2019), and upon treatment with apoptosis-inducing stressors, JMY expression is increased (Coutts et al., 2007; Coutts et al., 2009; Zuchero et al., 2012). Conclusions on JMY’s localization were largely drawn from fluorescence microscopy experiments observing overexpressed JMY constructs, where tagged JMY was mostly cytoplasmic before becoming concentrated in the nucleus in response to long-term exposures to stress. In contrast, immunofluorescence microscopy experiments visualizing endogenous JMY showed that JMY was primarily nuclear at steady state in multiple cell types (Coutts et al., 2007; Zuchero et al., 2009). Consistent with the latter observations, endogenous JMY also appeared to be nuclear in U2OS cells under normal growth conditions, and changes in its cytosolic and nuclear localization were G-protein regulated (King et al., 2021). In our current experiments, endogenous JMY exhibited a prominently nuclear localization in U2OS cells at steady state, while GFP-JMY maintained a mostly cytosolic presence (Figure 1A). These differences observed in the nucleo-cytoplasmic distribution of endogenous compared to tagged JMY imply that GFP-JMY is more useful for studying the cytosolic rather than nuclear activities of JMY.

To more closely evaluate the levels and localization of endogenous JMY in response to DNA damage, we exposed U2OS cells to etoposide for up to 8h before fixing and staining them to visualize endogenous JMY, F-actin, and DNA. We then quantified the whole cell JMY fluorescence for individual cells as well as the intensity of JMY in the nucleus and cytosol. Similar to previous findings of tagged JMY (Coutts et al., 2009; Zuchero et al., 2012), whole cell JMY fluorescence increased significantly by 6h of etoposide treatment (Figure 1C and S1; magenta stars). An assessment of JMY intensity in the cytosol and nucleus revealed a 3:7 ratio of cytosolic to nuclear JMY under basal conditions (Figure 1C and S1). By comparison, cytosolic JMY intensity increased significantly during 4-8h of etoposide exposure (Figure 1C; magenta stars), whereas nuclear JMY intensity was equivalent regardless of etoposide (Figure 1C and S1). Together, these results show that endogenous JMY normally exhibits a primarily nuclear localization and, upon introduction of an apoptosis-inducing stressor, JMY expression specifically increases in the cytosol while remaining high in the nucleus.

We hypothesized that the formation of juxtanuclear JMY puncta in response to etoposide was responsible for the increased cytosolic JMY intensity found in the whole cells. To test this, we assessed the levels and distribution of JMY in cells that contained clusters of punctate JMY compared to those that did not. When the total cell population was split into JMY puncta-positive and puncta-negative cells, the puncta-positive cells exhibited significantly higher whole cell JMY fluorescence at all timepoints compared to the puncta-negative cells (Figure 1C; left panel). When evaluating cytosolic JMY intensity specifically, the puncta-positive cells always showed >2-fold higher cytosolic JMY intensity compared to puncta-negative cells (Figure 1C; middle panel). This was in contrast to the nuclear JMY intensity, where the puncta-positive and puncta-negative populations displayed statistically indistinguishable levels of nuclear JMY regardless of the duration of etoposide treatment (Figure 1C and S1). This difference is further emphasized when comparing the ratio of cytosolic to nuclear JMY intensity in the two populations, where the puncta-negative cells maintained a 3:7 ratio of cytosolic to nuclear JMY intensity irrespective of etoposide treatment, while the puncta-positive cells displayed a nearly 1:1 ratio of cytosolic to nuclear JMY intensity after etoposide exposure (Figure S1). Thus, in response to DNA damage, cells undergo an overall increase in JMY expression due to the formation of a juxtanuclear cluster of JMY puncta.

Given that the best understood molecular behavior of JMY resides in its ability to promote actin assembly, we next investigated the timing of JMY puncta formation in relation to the reorganization of F-actin surrounding the JMY puncta. To visualize this process live, we utilized U2OS cells stably expressing GFP-JMY and transiently expressing the F-actin probe Lifeact-mCherry. The cells were treated with etoposide for 3h, and then imaged for an additional 1.5h. Selected frames taken from Video S1 show that the number and intensity of punctate GFP-JMY structures increased over time (Figure 1D). By comparison, during the same period, the cell began to round, but F-actin was not intensely concentrated in the juxtanuclear region until GFP-JMY levels reached an apparent threshold (Figure 1D). Fluorescence intensity profiles over time for two cytosolic areas within the cellular region containing punctate GFP-JMY structures showed GFP-JMY intensity increasing steadily and plateauing between 230 and 260 minutes, while Lifeact-mCherry intensity remained steady initially and then increased beginning at 230 minutes before achieving its maximum values between 240 and 260 minutes (Figure 1E; i-ii). In contrast, a cytosolic region lacking GFP-JMY puncta showed a decrease in Lifeact-mCherry intensity after 230 minutes (Figure 1E; iii), and a nuclear region did not display any dramatic changes in JMY or F-actin intensity over time (Figure 1E, nuclear). These results indicate that increased incorporation of JMY into a cluster of cytosolic puncta precedes the formation of a juxtanuclear F-actin-rich territory.

### Apoptotic F-actin-rich territories contain the JMY/WHAMM subfamily of Arp2/3-activating nucleation factors

JMY, its closest homolog WHAMM, and their downstream branched actin nucleator, the Arp2/3 complex, are all required for efficient intrinsic apoptosis (King et al., 2021). Therefore, we next studied the localization of these and other actin nucleation and branching factors in relation to the F-actin-rich territories. To do this, we treated U2OS cells with etoposide for 6h and visualized endogenous JMY or GFP-JMY relative to F-actin, the Arp2/3 complex, and the branching regulator Cortactin. Two Arp2/3 complex subunits, Arp3 and ArpC2, along with Cortactin, localized to the JMY-containing territories (Figure 2A and S2). Fluorescence intensity plot profiles of Arp2/3 subunit localizations showed increased intensity throughout the entire JMY- and F-actin regions compared to adjacent cytosolic areas (Figure 2B), and magnifications indicated that the Arp2/3 complex often overlapped with intense F-actin structures interspersed throughout the territories (Figure 2A and 2B). In contrast, fluorescence profiles of Cortactin, which binds actin and the Arp2/3 complex to modulate branchpoint stability (Schnoor et al., 2018), typically showed increased Cortactin intensity at peripheral actin filaments surrounding interior punctate JMY structures (Figure S2). Therefore, the apoptotic F-actin-rich regions contain the canonical components of the branched actin assembly machinery, with the Arp2/3 complex localizing throughout the territory and Cortactin found mainly at the territory perimeter.

**Figure 2.**
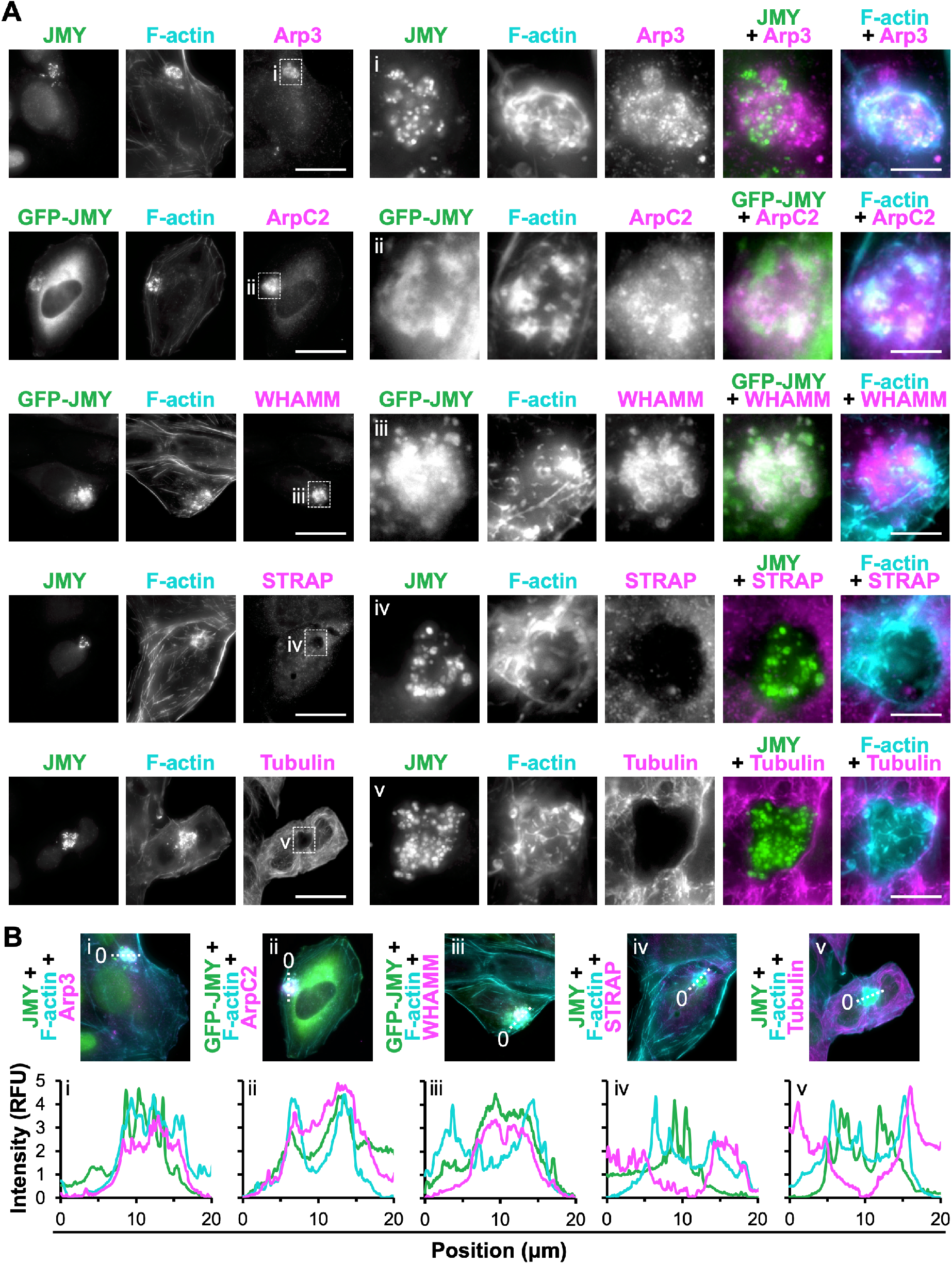
A subset of actin nucleation and branching factors is present within apoptotic F-actin-rich territories. **(A)** U2OS cells or U2OS cells expressing GFP-JMY (green) were treated with etoposide for 6h, fixed, and stained with antibodies to JMY (green), antibodies to *-* Arp3, ArpC2, WHAMM, STRAP, or tubulin (magenta), and with phalloidin (F-actin; cyan). Magnifications (i-iii) depict clusters of JMY and other nucleation factors in F-actin territories. Magnifications (iv-v) depict territories that exclude STRAP or tubulin. Scale bars, 25μm, 5μm. **(B)** 20μm lines were drawn through the images in (A) using ImageJ to measure the pixel intensity profiles. RFU = relative fluorescence units. See Figure S2 for nucleation factors that were not enriched in F-actin territories.

Genetic inactivation of each WASP-family Arp2/3 activator previously revealed that only cells lacking JMY or WHAMM have substantial deficiencies in their intrinsic apoptosis responses (King et al., 2021). To characterize which, if any, WASP-family proteins other than JMY associate with the apoptotic F-actin territories, we treated U2OS cells with etoposide and assessed the localization of WHAMM and representatives of other subgroups within the WASP family. Endogenous and tagged WHAMM, but not N-WASP, WAVE2, or WASH, exhibited strong localization to the JMY- and F-actin-rich regions (Figure 2A and S2). Magnified insets highlighted the occasional colocalization of JMY, WHAMM, and F-actin (Figure 2A and S2), and fluorescence intensity plot profiles showed increased WHAMM intensity in close proximity to JMY within F-actin territories (Figure 2B and S2). These results demonstrate that the cytosolic F-actin-rich territories which form during the intrinsic apoptotic response contain a specific subset of nucleation factors, namely the Arp2/3 complex, JMY, and WHAMM.

The different localizations and functions of WASP-family members are controlled via interactions with regulatory complexes, small G-proteins, phospholipids, and microtubules, but little is known about how upstream factors regulate the localization or actin assembly activity of JMY or WHAMM under apoptotic conditions. In certain settings, the stress-response protein STRAP may inhibit JMY-mediated nucleation and interrupt its cytosolic functions while facilitating its role as a co-activator of transcription in the nucleus (Demonacos et al., 2001; Hu and Mullins, 2019). So we next examined the localization of STRAP relative to JMY and found that, in stark contrast to the enhanced localization of JMY and the Arp2/3 complex in the apoptotic F-actin territories, STRAP was excluded from these regions (Figure 2A and 2B). For WHAMM, binding to microtubules inhibits its nucleation-promoting activity (Campellone et al., 2008; Shen et al., 2012). Notably, microtubules were also not found within the JMY-containing F-actin territories (Figure 2A), as magnifications and fluorescence intensity plot profiles showed a clear absence of tubulin staining in such regions (Figure 2A and 2B). Hence, the apoptotic F-actin-rich territories not only consist of specific components of the branched actin nucleation machinery, but also act like compartments that exclude inhibitors of both JMY- and WHAMM-mediated actin polymerization.

### The core apoptosome components cytochrome *c* and Apaf-1 are enriched within juxtanuclear F-actin-rich territories

Intrinsic pathways of apoptosis are characterized by the export of apoptogenic proteins including cyto *c* from mitochondria to the cytosol, leading to the initiation of caspase cleavage cascades (Tait and Green, 2013; Bock and Tait, 2020). Previous work showed that punctate JMY structures can overlap with mitochondria-independent cyto *c* and that cytosolic cyto c puncta are not detectable in JMY knockout cells (King et al., 2021), so we further evaluated the relationship of cyto *c* to the JMY- and Arp2/3-containing actin-rich regions. Consistent with earlier observations, many bright JMY puncta colocalized with cytosolic cyto *c* puncta, and these structures were clustered within F-actin territories (Figure 3A). Such cyto *c* puncta formed in close proximity to Arp2/3 complex and WHAMM structures, but with less overlap (Figure 3A).

**Figure 3.**
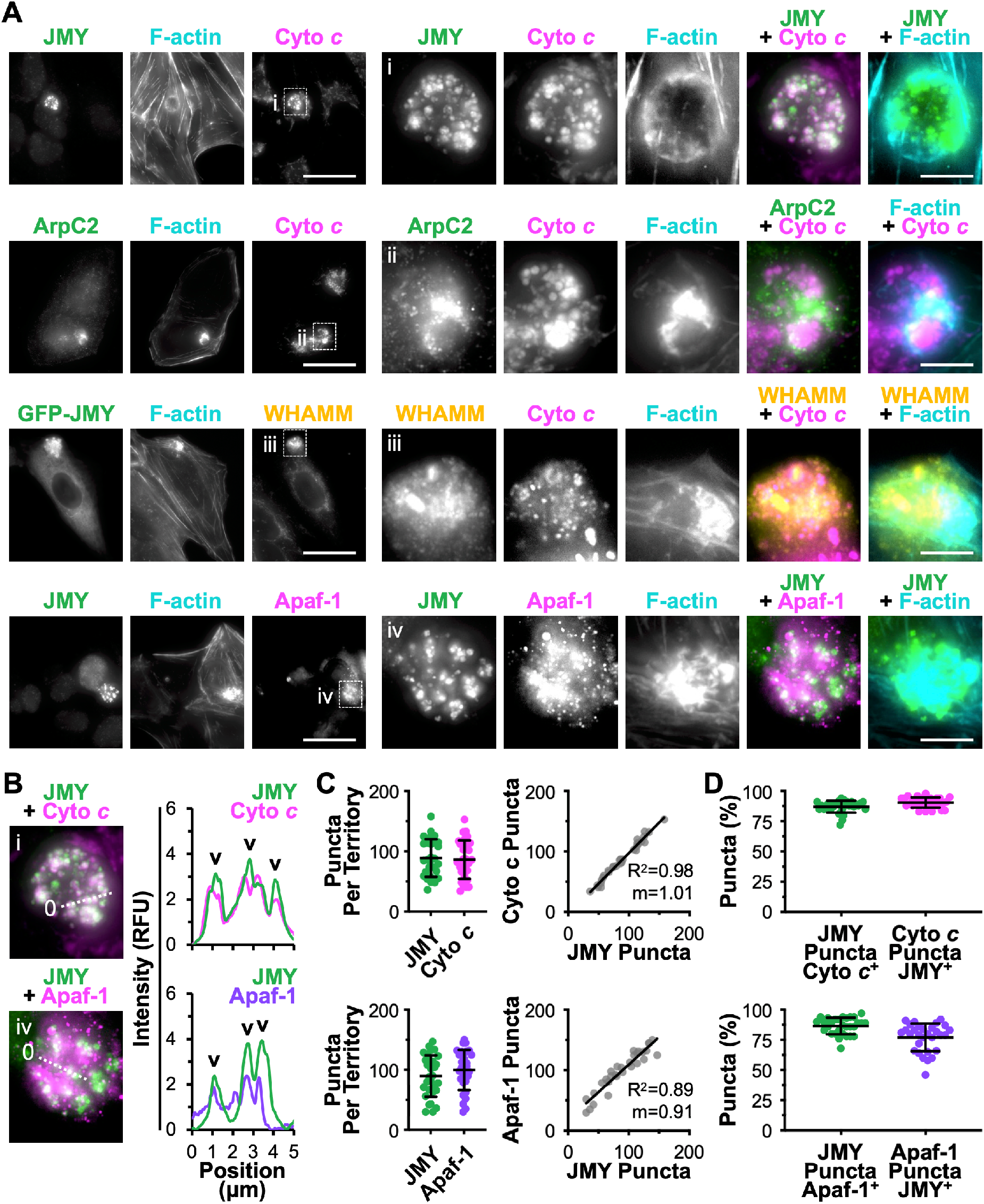
The apoptosome components cytochrome *c* and Apaf-1 are clustered within F-actin-rich territories. **(A)** U2OS cells or U2OS cells expressing GFP-JMY (green) were treated with etoposide for 6h, fixed, and stained with antibodies to JMY (green), cyto *c* or Apaf-1 (magenta), and/or WHAMM (yellow), and with phalloidin (F-actin; cyan). Magnifications (i-iv) depict JMY, WHAMM, cyto *c*, and Apaf-1 within F-actin territories. Scale bars, 25μm, 5μm. **(B)** 5μm lines were drawn through the magnified images in (i,iv) using ImageJ to measure the pixel intensity profiles. Arrowheads highlight examples of cytosolic cyto *c* or Apaf-1 puncta that *-* overlap with JMY puncta. **(C)** For quantification of (A), the number of JMY and cyto *c* or Apaf-1 puncta were counted per territory. Each point represents 1 cell (n = 30). The number of cyto *c* or Apaf-1 puncta per territory was plotted against the number of JMY puncta per territory. The slopes (m) in the linear trendline equations for cyto *c* (Y = 1.01X – 3.61) and Apaf-1 (Y = 0.91X + 18.02) were significantly non-zero (p<0.001). **(D)** Quantifications show the % of JMY puncta per territory that overlapped with cyto *c* or Apaf-1 puncta (cyto *c*-or Apaf-1-positive) and the % of cyto *c* or Apaf-1 puncta per territory that overlapped with JMY puncta (JMY-positive). See Figure S3 for other factors that were not enriched in F-actin territories.

After cyto *c* is released into the cytosol, it interacts with the scaffolding protein Apaf-1 and triggers Apaf-1 oligomerization into heptameric complexes called apoptosomes, which serve as platforms for the multimerization and proteolytic processing of caspases (Bratton and Salvesen, 2010; Yuan and Akey, 2013; Dorstyn et al., 2018). To determine if Apaf-1-containing apoptosomes might be forming within the F-actin territories, we stained cells with Apaf-1 antibodies. Apaf-1 puncta were indeed enriched within the territories and appeared to overlap frequently with JMY structures (Figure 3A). Fluorescence intensity plot profiles showed colocalization between JMY and cyto *c* puncta as well as between JMY and several larger Apaf-1 puncta (Figure 3B). Therefore, the cytosolic JMY-containing F-actin-rich areas encompass both of the core apoptosome proteins.

To better understand if JMY influences apoptosome formation within the territories, we characterized the abundance of apoptosome components in relation to JMY. Quantification of the number of individual puncta within apoptotic F-actin-rich territories gave median values of 89 JMY puncta and 86 cyto *c* puncta per territory, with a wide range of puncta present within individual territories (Figure 3C). Moreover, the number of cyto *c* puncta showed a positive correlation with the number of JMY puncta (Figure 3C). When assessing their spatial overlap, 87% of JMY puncta were cyto *c*-positive and 90% of cyto *c* puncta were JMY-positive (Figure 3D), meaning not only are equivalent numbers of these punctate structures forming within F-actin-rich territories, but the majority of JMY and cyto *c* puncta are at least partially localizing together. Assessments of JMY and Apaf-1 structures within F-actin territories revealed similar phenotypes, including median values of 89 JMY puncta and 100 Apaf-1 puncta per territory and a positive correlation between the numbers of JMY and Apaf-1 puncta (Figure 3C). 87% of JMY puncta were Apaf-1-positive while 77% of Apaf-1 puncta were JMY-positive (Figure 3D), indicating that the majority of JMY and Apaf-1 structures also localize together, although not to the same extent as JMY and cyto *c*. Collectively, these results suggest that individual F-actin-rich territories harbor relatively stoichiometric amounts of JMY, cyto *c*, and Apaf-1 structures.

For determining the specificity with which such pro-apoptotic factors accumulate in the territories, we also examined the localization of other cellular components by visualizing p53, mitochondria, or the Golgi in relation to JMY and F-actin. Consistent with a primarily nuclear function for p53 during apoptosis, it was not enriched within cytosolic JMY-associated F-actin territories (Figure S3). Additionally, while cyto *c* is the most-well studied apoptogenic protein that is exported from permeabilized mitochondria during apoptosis, another mitochondrial protein, AIF, was not present within the territories (Figure S3). WHAMM and JMY can promote vesicle trafficking to and from the Golgi, but a Golgi membrane marker, GM130, was not found within the F-actin regions either (Figure S3). These findings further support the idea that F-actin-rich territories function in compartmentalizing specific pro-apoptotic factors in close proximity to one another to allow for the efficient formation of apoptosomes.

### Apoptotic F-actin-rich territories concentrate active executioner caspases

Apoptosomes multimerize and activate initiator and executioner caspases, the latter of which eventually cleave multiple proteins in the cytosol and nucleus to drive apoptotic cell death (Dorstyn et al., 2018; Bock and Tait, 2020). While structural and biochemical studies have shown apoptosomes to be important for starting caspase cleavage cascades by facilitating interactions between Apaf-1 and initiator procaspase-9 and promoting its proteolytic activity (Li et al., 2017; Su et al., 2017; Wang et al., 2017), the spatiotemporal mechanisms underlying caspase recruitment to, and activation by, apoptosomes in cells are not well understood. We previously demonstrated that JMY knockout fibroblasts have defects in the cleavage of initiator caspase-9 and executioner caspase-3, and that cleaved caspase-3 can be found in association with JMY- and F-actin-rich areas of cells (King et al., 2021). So to evaluate the localization of initiator caspase-9 and executioner caspase-3 in relation to apoptotic F-actin-rich territories, we stained etoposide-treated cells with antibodies to total caspase-9 (TCasp-9) and total caspase-3 (TCasp-3), which recognize both their inactive full length and active cleaved forms. Both TCasp-9 and TCasp-3 were present in the JMY-containing F-actin regions (Figure S4).

To next determine whether the apoptosome-containing territories were enriched for active caspases, cells were stained with antibodies that specifically recognize active cleaved caspase-3 (CCasp-3). CCasp-3 fluorescence was usually present entirely within F-actin-rich territories (Figure 4A, 4B, and S4). CCasp-3 staining often coincided with JMY and WHAMM structures, and also associated with intense clusters of punctate cyto *c* and Apaf-1 (Figure 4A), indicating that JMY, WHAMM, apoptosomes, initiator caspases, and active executioner caspases altogether localize within F-actin-rich territories.

**Figure 4.**
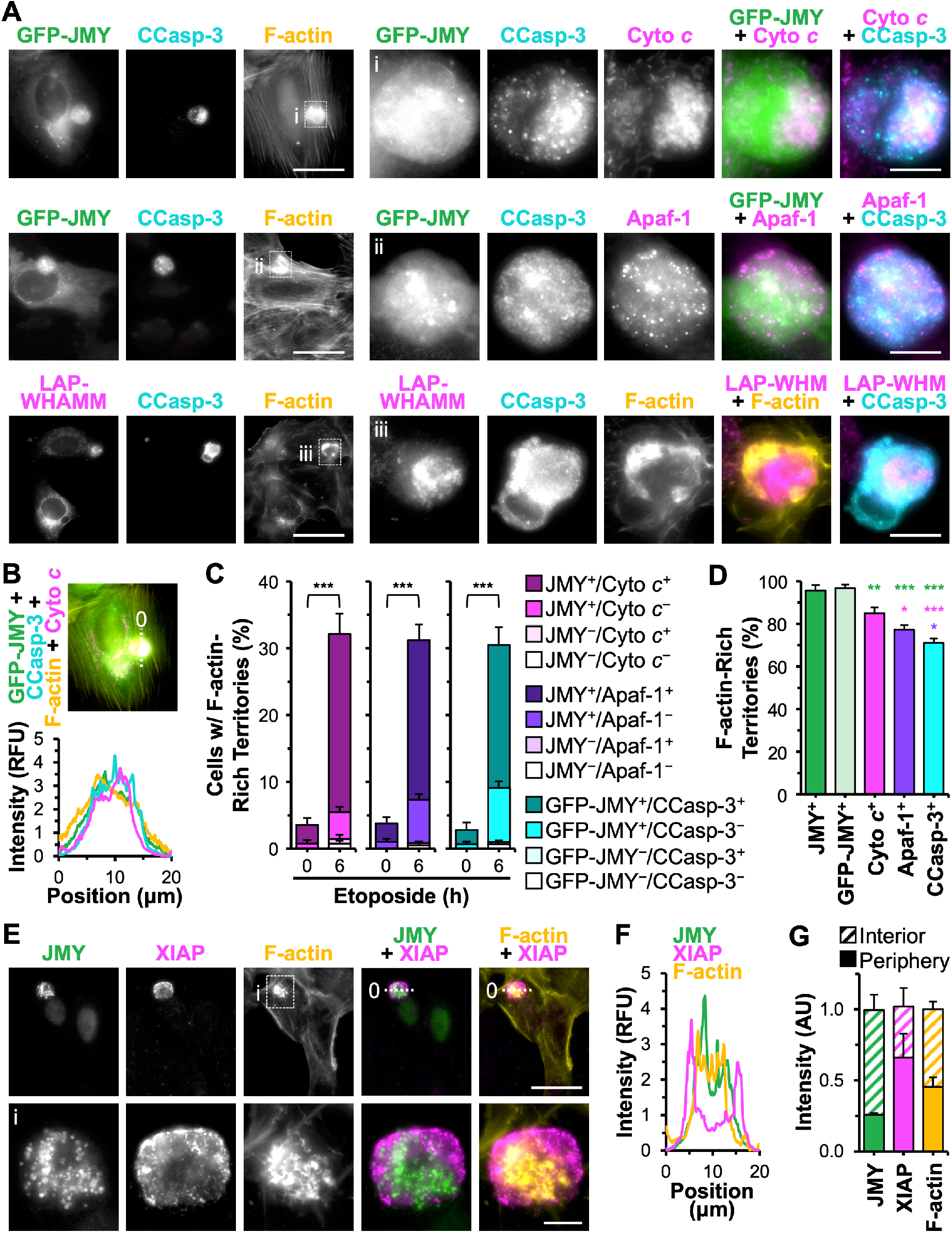
Active executioner caspase-3 is concentrated within apoptosome-containing F-actin territories. **(A)** U2OS cells expressing GFP-JMY (green) or transfected with a plasmid encoding LAP-WHAMM (magenta) were treated with etoposide for 6h, fixed, and stained with *-* an antibody that recognizes cleaved caspase-3 at Asp175 (CCasp-3; cyan), antibodies against cyto *c* or Apaf-1 (magenta), and with phalloidin (F-actin; yellow). Magnifications (i-iii) depict GFP-JMY, cyto *c*, Apaf-1, cleaved caspase-3, and LAP-WHAMM within F-actin territories. Scale bars, 25μm, 5μm. **(B)** A 20μm line was drawn through an image from (A) using ImageJ to measure the pixel intensity profiles for GFP-JMY, cleaved caspase-3, F-actin, and cyto *c*. **(C-D)** U2OS cells or U2OS cells expressing GFP-JMY were untreated or treated with etoposide, fixed, and stained with antibodies against JMY and cyto *c*, JMY and Apaf-1, or cleaved caspase-3, all in conjunction with phalloidin. The % of cells harboring F-actin-rich territories containing JMY, GFP-JMY, cyto *c*, or Apaf-1 puncta, or cleaved caspase-3 clusters was quantified. In (C), the fraction of F-actin-rich territories that were doubly-positive, singly-positive, or doubly-negative are shown. Each bar represents the mean±SD from 3 experiments (n = 504-638 cells per bar). ***p<0.001 (t-tests). In (D), each bar represents the mean±SD from 3 experiments (n = 182-195 territories per bar). Green, magenta, and purple significance stars refer to comparisons to the JMY^+^ bar, Cyto *c*^+^ bar, and Apaf-1^+^ bar respectively. **(E)** U2OS cells were treated with etoposide, fixed, and stained with antibodies to JMY (green) and XIAP (magenta), and with phalloidin (F-actin; yellow). Magnifications (i) depict F-actin territories surrounded by XIAP. **(F)** A 20μm line was drawn through the image in (E) using ImageJ to measure the pixel intensity profiles for JMY, XIAP, and F-actin. **(G)** ImageJ was used to measure the JMY, XIAP, and F-actin fluorescence intensities for the whole territory as well as the interior and periphery portions (n = 17 territories). *p<0.05; **p<0.01; ***p<0.001 (ANOVA, Tukey post-hoc tests).

Intrinsic pathways of apoptosis have been described as stepwise progressions starting with the release of cyto *c* from mitochondria, which triggers the assembly of the Apaf-1 apoptosome, allowing for the activation of initiator caspases that then activate executioner caspases. We next wanted to place JMY and F-actin more precisely within this pathway by measuring the frequency with which F-actin-rich territories harbored JMY puncta, apoptosome components, and active caspases. Similar to the trends shown for JMY puncta formation following DNA damage (Figure 1B), F-actin-rich territories were present in >30% of cells after 6h of etoposide treatment (Figure 4C). Furthermore, the majority of F-actin territories contained a pairwise combination of JMY with cyto *c*, Apaf-1, or CCasp-3 staining (Figure 4C), as ∼25% of cells included F-actin-rich regions that were positive for both JMY and apoptosome components while ∼20% were positive for both GFP-JMY and CCasp-3 (Figure 4C). In contrast, only 5-10% of cells had F-actin territories containing JMY puncta without apoptosome components or active executioner caspases (Figure 4C). F-actin-rich areas that lacked JMY, cyto *c*, Apaf-1, and CCasp-3 staining were even rarer (Figure 4C). Interestingly, when quantifying the proportion of F-actin territories that stained for JMY, apoptosome proteins, or active executioner caspases, >96% of F-actin territories were JMY puncta-positive, 85% were cyto *c*-positive, 77% were Apaf-1-positive, and 71% were CCasp-3-positive (Figure 4D). These frequencies with which F-actin-rich territories incorporate JMY puncta, apoptosomes, and executioner caspases are consistent with the conclusion that a stepwise pathway from mitochondrial permeabilization to apoptosome biogenesis to caspase activation all takes place within this discrete domain of the cytosol.

Apoptosis is tightly controlled by both positive and negative regulators, and members of the inhibitor of apoptosis protein (IAP) family prevent apoptosis by blocking caspase activation or activity (Hrdinka and Yabal, 2019). One potent caspase inhibitor is XIAP, which binds directly to initiator caspase-9 and executioner caspases-3 and -7, thereby blocking their proteolytic activities (Deveraux and Reed, 1999; Scott et al., 2005; Eckelman et al., 2006). We next evaluated the spatial relationship of XIAP to JMY and F-actin. XIAP was surprisingly recruited to JMY-containing F-actin-rich territories (Figure 4E). However, fluorescence intensity plot profiles showed XIAP intensity peaking at the peripheral boundaries of the territory rather than in association with the interior JMY structures (Figure 4E and 4F). Measurements of JMY and XIAP levels at the inner versus outer portions of multiple territories revealed that JMY was most abundant in internal regions of the territories whereas XIAP was significantly enriched at their perimeters (Figure 4G). These results indicate that the apoptotic F-actin-rich territories not only compartmentalize apoptosomes, initiator caspases, and executioner caspases, but also feature a specialized peripheral positioning of the caspase inhibitor XIAP.

### Localized JMY-mediated Arp2/3 activation and actin polymerization are crucial for driving executioner caspase activation

While previous work showed that JMY requires its actin- and Arp2/3-binding domains in order to enable efficient caspase-3 cleavage (King et al., 2021), the degree to which actin assembly specifically within territories impacts caspase-3 processing is unknown. JMY activates the Arp2/3 complex using a C-terminal domain comprising 3 actin-binding WASP-homology-2 (W) motifs and an Arp2/3-binding Connector Acidic (CA) segment, and can also use its WWW region to nucleate actin directly (Zuchero et al., 2009). JMY knockout (JMY^KO^) fibroblasts are deficient in caspase-3 activation (King et al., 2021), so we utilized 3 GFP-tagged JMY derivatives for performing rescue experiments: wild type JMY (JMY^WT^); a JMY mutant missing all 3 W motifs (JMY^ΔWWW^) that can neither nucleate actin directly nor cooperate with the Arp2/3 complex; and a JMY mutant lacking the CA segment (JMY^ΔCA^) that cannot activate Arp2/3 but retains its actin nucleating WWW region. We introduced the GFP-JMY plasmids, or a vector control, into the JMY^KO^ cells, exposed them to etoposide, and stained them to visualize active CCasp-3 and F-actin (Figure 5A). Similar to previous observations (King et al., 2021), just ∼5% of vector- or JMY mutant-transfected JMY^KO^ cells stained positive for active CCasp-3 (Figure 5A and 5B), while >15% of cells transfected with the full-length JMY construct were CCasp-3-positive (Figure 5A and 5B), indicating that only wild type JMY could promote executioner caspase activation in JMY^KO^ cells.

**Figure 5.**
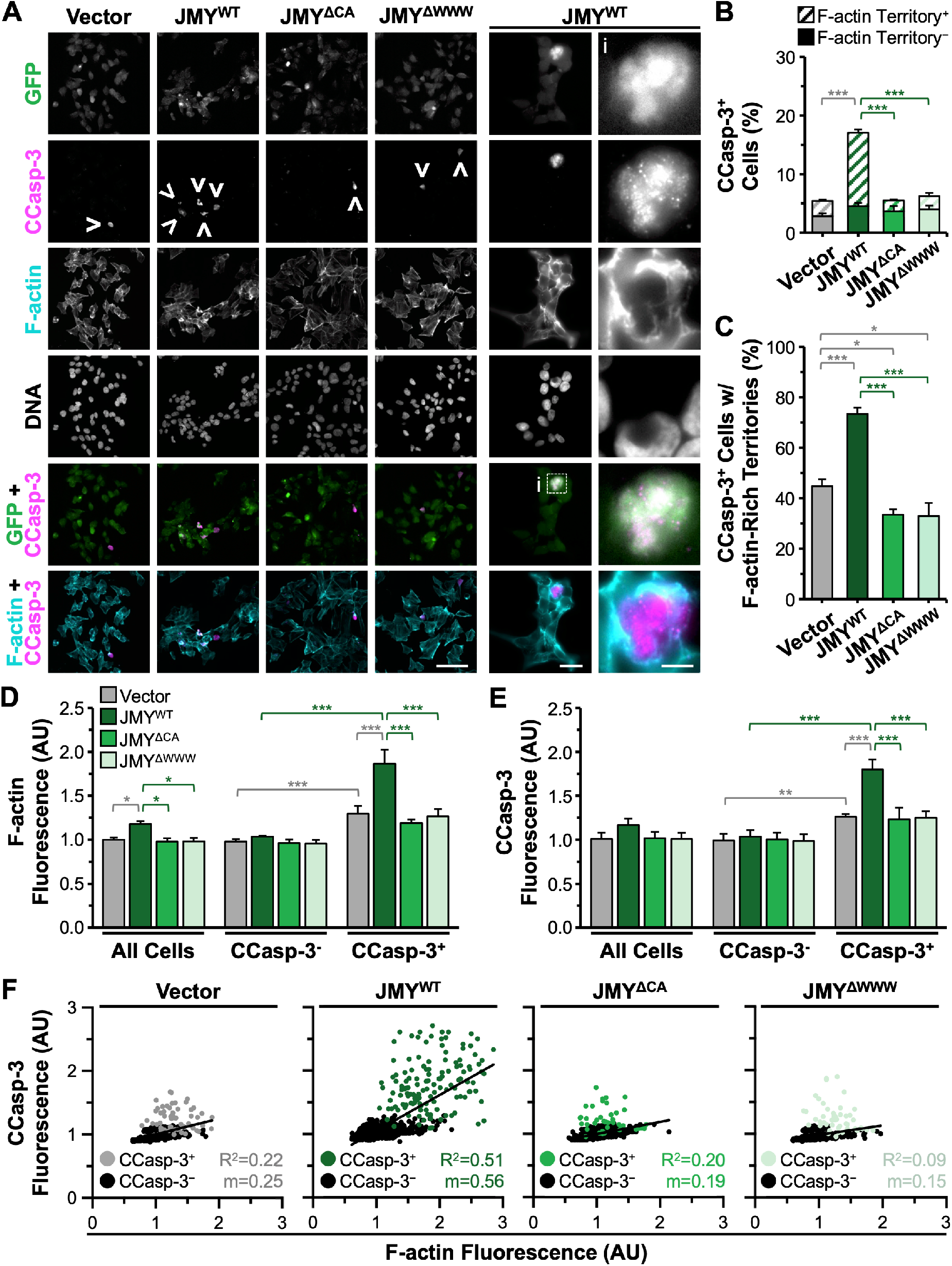
The localized activation of caspase-3 is controlled by JMY-dependent actin polymerization. **(A)** JMY^KO^ cells harboring plasmids encoding GFP (vector) or GFP-tagged JMY constructs (JMY^WT^, JMY^ΔCA^, JMY^ΔWWW^; green) were treated with etoposide for 5h, fixed, *-* and stained with an antibody to visualize cleaved caspase-3 (CCasp-3; magenta), phalloidin (F-actin; cyan), and DAPI to detect DNA. Arrowheads highlight examples of CCasp-3 staining, and magnifications (i) depict GFP-JMY^WT^ and CCasp-3 surrounded by F-actin. Scale bars, 100μm, 25μm, 5μm. **(B)** The % of cells with cleaved caspase-3 staining was quantified and displayed as the fraction of CCasp-3-positive cells that were (Territory^+^) or were not (Territory^-^) associated with F-actin. Each bar represents the mean±SD from 3 experiments (n = 3,447-6,083 cells per bar). Gray significance stars refer to comparisons to the vector bar and green significance stars are comparisons to the JMY^WT^ bar. **(C)** The % of cells with CCasp-3 staining associated with F-actin was quantified. Each bar is the mean±SD from 3 experiments (n = 1,038 cells for WT and 190-230 for vector and mutant samples). **(D-E)** Whole cell fluorescence values for F-actin and CCasp-3 were measured and the mean value for the vector sample was set to 1. Each bar is the mean±SD from 3 experiments (n = 592-816 cells per sample). **(F)** Whole cell fluorescence values for F-actin and CCasp-3 were plotted against one another. Each point represents 1 cell (n = 592-816 cells per sample) where gray/green points are CCasp-3-positive cells and black points are CCasp-3-negative cells. The slopes (m) in the linear trendline regression equations for the total cell populations (vector: Y = 0.25X + 0.75; JMY^WT^: Y = 0.56X + 0.49; JMY^ΔCA^: Y = 0.19X + 0.81; JMY^ΔWWW^: Y = 0.15X + 0.84) were significantly non-zero (p<0.001). *p<0.05; **p<0.01; ***p<0.001 (ANOVA, Tukey post-hoc tests).

To explore the precise relationship between activated caspase-3 and F-actin territories, the proportion of CCasp-3-positive cells that had active caspase staining localized within the F-actin-rich regions was quantified. Out of the infrequent CCasp-3-positive cells observed in the vector-transfected JMY^KO^ sample, less than half were associated with F-actin-rich territories (Figure 5B and 5C). Compared to CCasp-3-positive cells that had strong active caspase staining clustered within F-actin-rich regions, those that were independent of F-actin displayed weaker CCasp-3 staining that was more diffuse throughout the cell. In contrast, for samples transfected with the wild type JMY construct, 75% of the active CCasp-3-positive cells had their staining encompassed within F-actin-rich territories (Figure 5B and 5C). For both JMY mutants, the frequency of active caspase staining that associated with F-actin-rich territories slightly decreased relative to vector-transfected cells (Figure 5C). These results show that the actin polymerization capacity of JMY is required for localized caspase activation specifically within F-actin-rich territories.

We next asked if there was any difference in the amount of F-actin or cleaved caspase-3 in the absence or presence of the different JMY derivatives by quantifying the F-actin (Figure 5D) and CCasp-3 (Figure 5E) fluorescence intensities for individual cells. Mean F-actin fluorescence values were higher in the cells transfected with wild type JMY compared to the vector- or mutant-transfected JMY^KO^ cells (Figure 5D). Moreover, when total cell populations were split into CCasp-3-positive and -negative cells, the CCasp-3-positive cells showed greater F-actin fluorescence compared to the negative cells, and the JMY^WT^ re-expressing cells contained significantly more F-actin than the vector-transfected JMY^KO^ cells (Figure 5D). These results indicate that the increased whole cell F-actin content observed within the JMY^WT^ sample can be attributed to the apoptotic territories. Additionally, within the CCasp-3-positive population, JMY^WT^ re-expressing cells also displayed significantly higher CCasp-3 fluorescence compared to the vector-expressing cells (Figure 5E). The JMY^ΔWWW^ and JMY^ΔCA^ samples behaved identically to vector-transfected cells in assays measuring both F-actin and CCasp-3 levels (Figure 5D and 5E). Thus, JMY-mediated actin assembly drives a more frequent and more efficient activation of executioner caspases when they are confined within territories.

To directly relate F-actin abundance to caspase-3 activation, we plotted these two parameters against one another on a cell-by-cell basis. Comparisons of CCasp-3 to F-actin fluorescence in individual cells showed that those transfected with wild type JMY exhibited a stronger positive correlation between the intensity of CCasp-3 and F-actin fluorescence than did vector- or mutant-transfected cells (Figure 5F). These results show that the amount of actin polymerized via JMY has a direct relationship with the potency of caspase-3 activation in defined cytosolic territories.

### JMY and WHAMM both contribute to F-actin-rich territory assembly and clustering-mediated caspase activation

Although JMY is crucial for assembling apoptotic F-actin territories in which caspase-3 is activated, some JMY-deficient cells still contain active caspase-3 in association with actin (Figure 5B), so we next wanted to identify the molecular basis for forming these cytoskeletal compartments in the presence and absence of JMY. Based on our findings that efficient apoptosis requires JMY and WHAMM (King et al., 2021), and that the territories contain these two specific nucleation factors (Figure 2 and S2), we evaluated the relative contributions of JMY and WHAMM to the assembly of F-actin-rich territories and caspase activation. We employed two independent JMY KO fibroblast cell lines, two independent WHAMM KO lines, and two WHAMM/JMY double knockout (DKO) lines, exposed them to etoposide, and stained them to visualize active CCasp-3 and F-actin. Similar to previous observations (King et al., 2021), ∼40% of parental (HAP1, eHAP) cells stained positive for active CCasp-3, whereas only ∼20% of WHAMM^KO^ and <10% of the JMY^KO^ or WHAMM/JMY^DKO^ cells were positive for CCasp-3 staining (Figure 6A and 6B), reaffirming that wild type cells can execute an efficient cell death program whereas JMY^KO^ or WHAMM^KO^ cells have significant deficiencies in executioner caspase activation.

**Figure 6.**
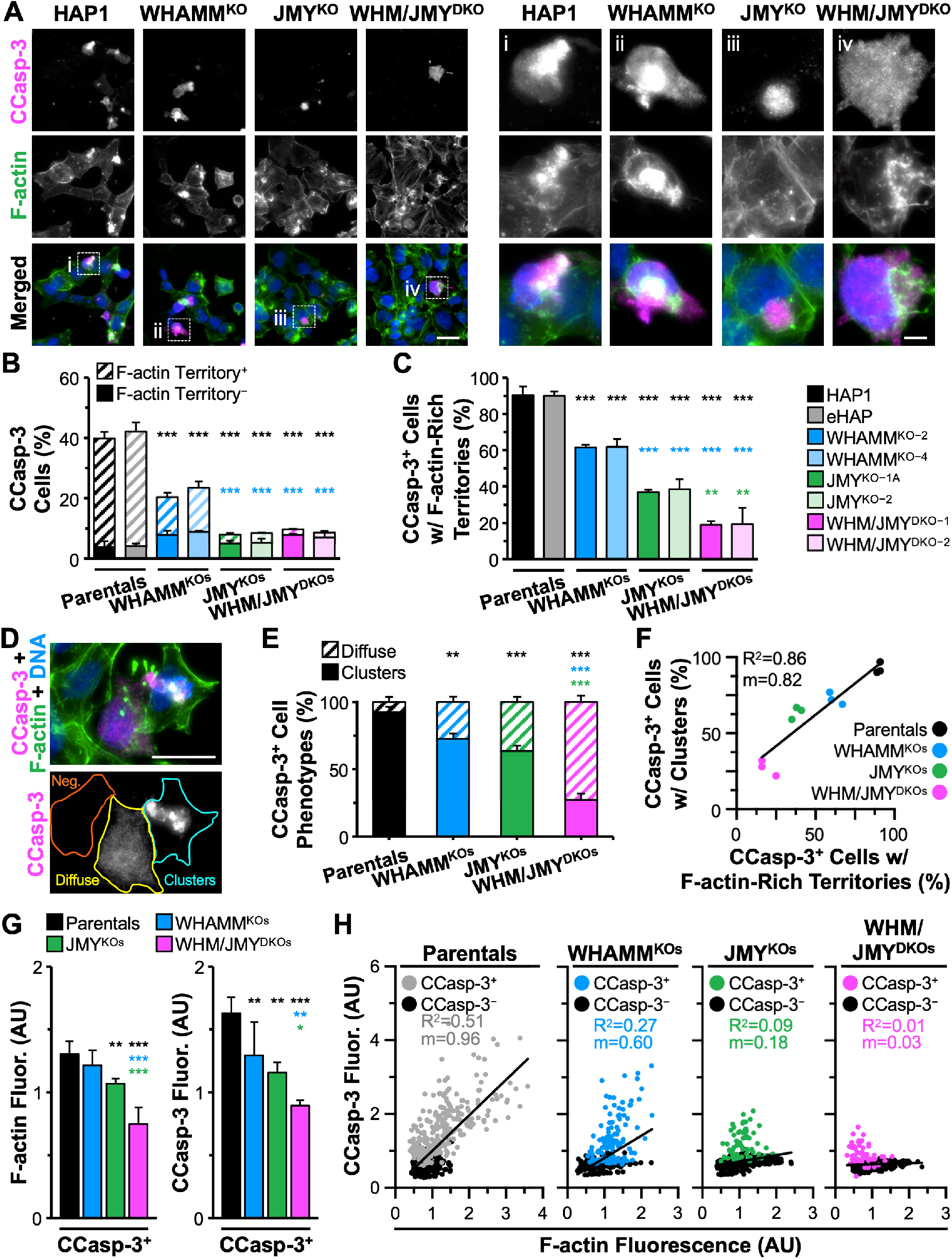
The amount of F-actin encompassing the caspase activation clusters is enhanced by WHAMM. **(A)** Parental (HAP1, eHAP), WHAMM^KO^ (WHAMM^KO-2^, WHAMM^KO-4^), JMY^KO^ (JMY^KO-1A^, JMY^KO-2^), and WHAMM/JMY^DKO^ (WHM/JMY^DKO-1^, WHM/JMY^DKO-2^) cells were *-* treated with 5μM etoposide for 6h before fixing and staining with an antibody to visualize cleaved caspase-3 (CCasp-3; magenta), phalloidin (F-actin; green), and DAPI (blue). Magnifications (i-iv) show examples of CCasp-3 staining. Scale bars, 25μm, 5μm. **(B)** The % of cells with CCasp-3 staining was quantified and displayed as the fraction of CCasp-3-positive cells that were (Territory^+^) or were not (Territory^-^) associated with F-actin-rich territories. Each bar represents the mean±SD from 3 experiments (n = 246-438 cells per bar). Black significance stars refer to comparisons to the parental bars and blue significance stars are comparisons to the WHAMM^KO^ bars. **(C)** The % of CCasp-3-positive cells with F-actin-rich territories is displayed. Each bar represents the mean±SD from 3 experiments (n = 104-106 cells for parental bars; 66-73 cells for WHAMM^KO^ bars; and 31-36 cells for JMY^KO^ or WHM/JMY^DKO^ bars). Black significance stars are comparisons to the parental bars, blue significance stars are to the WHAMM^KO^, and green significance stars are to the JMY^KO^. **(D)** A representative image of JMY^KO^ cells highlights cells that are CCasp-3-negative (orange) or CCasp-3-positive and diffuse (yellow) or in clusters (cyan). Scale bar, 25μm. **(E)** The % of CCasp-3-positive cells with clustered or diffuse phenotypes was quantified. Each bar represents the mean±SD from 3 experiments (n = 63-98 cells per bar). **(F)** The % of CCasp-3-positive cells with a clustered phenotype was plotted against the % of CCasp-3-positive cells with F-actin territories. Each point represents the mean from an individual experiment (n = 18-39 cells per point; 63-98 cells per sample). The slope (m) in the linear trendline equation (Y = 0.82X + 20.97) was significantly non-zero (p<0.001). **(G)** Whole cell fluorescence values for F-actin and CCasp-3 were measured in individual cells and normalized to the parental sample. Each bar is the mean±SD from 3 experiments (n = 607-866 cells per sample). Total and CCasp-3-negative cell data appear in Figure S5. **(H)** Whole cell fluorescence values for CCasp-3 and F-actin were plotted against one another. Each point represents 1 cell (n = 607-866 cells per plot) where non-black points are CCasp-3-positive and black points are CCasp-3-negative. The slopes (m) in the linear regression equations for parentals (Y = 0.96X + 0.04), WHAMM^KOs^ (Y = 0.60X + 0.20), and JMY^KOs^ (Y = 0.18X + 0.53) were significantly non-zero (p<0.001), while the slope for the WHAMM/JMY^DKO^ sample (Y = 0.03X + 0.61) was not (p=0.070). *p<0.05; **p<0.01; ***p<0.001 (ANOVA, Tukey post-hoc tests in B-D; Fisher’s exact test in E).

To determine the extent to which each cell line activated caspase-3 within F-actin-rich territories, we quantified the percentage of CCasp-3-positive cells that were also positive for F-actin. In >90% of CCasp-3-positive parental cells, caspase staining was found in F-actin-rich territories (Figure 6B and 6C). In contrast, ∼60% of WHAMM^KO^, <40% of JMY^KO^, and <20% of WHAMM/JMY^DKO^ cells had CCasp-3 staining in conjunction with actin filaments (Figure 6C). Not surprisingly, two caspase staining phenotypes for CCasp-3-positive cells were apparent – clustered or diffuse (Figure 6D) – and classifying cells in these two categories even without phalloidin staining was predictive of the genotype of the cells. Independent assessments of the fraction of CCasp-3-positive cells exhibiting each phenotype demonstrated that >90% of parental CCasp-3-positive cells displayed a clustered staining pattern, leaving <10% with a diffuse localization (Figure 6E). The proportion of diffuse CCasp-3-positive cells increased to >25% of WHAMM^KO^, >35% of JMY^KO^, and >70% of WHAMM/JMY^DKO^ CCasp-3-positive cells (Figure 6E). When reincorporating the F-actin staining data, the percentage of CCasp-3-positive cells showing a clustered phenotype positively correlated with the percentage of CCasp-3-positive cells containing F-actin territories (Figure 6F). Thus, JMY or WHAMM knockouts result in less frequent activation of caspase-3 and less frequent caspase clustering within F-actin-rich territories. While an individual WHAMM mutation results in partial deficits in the formation of CCasp-3-associated F-actin territories, and JMY inactivation causes a severe impairment, a WHAMM/JMY double mutation causes an even more extreme defect.

Finally, to define how deleting WHAMM and/or JMY influenced the amount of F-actin and active caspase-3 in apoptotic cells, we quantified the F-actin and CCasp-3 fluorescence within individual cells. Compared to the parental cell lines, F-actin fluorescence values were not significantly different upon deletion of WHAMM and/or JMY in the CCasp-3-negative population (Figure S5). However, in the CCasp-3-positive population, CCasp-3 and F-actin levels each displayed stepwise decreases when WHAMM, JMY, or both factors were deleted (Figure 6G). On a cell-by-cell basis, wild type cells showed a correlation between CCasp-3 and F-actin abundance, as well as a clear separation between non-apoptotic versus apoptotic caspase-positive populations (Figure 6H). In contrast, the WHAMM^KO^ and JMY^KO^ cells showed weaker correlations between CCasp-3 and F-actin fluorescence, with a more extreme phenotype observed in the JMY^KOs^ (Figure 6H). Notably, the WHAMM/JMY^DKOs^ did not show any correlation between CCasp-3 and F-actin fluorescence per cell and no obvious separation between the non-apoptotic and apoptotic populations (Figure 6H). These results demonstrate that while the loss of JMY alone inhibits the formation of intense F-actin-rich regions and the activation of caspase-3, these deficits are exacerbated upon the additional deletion of WHAMM, pointing to a functional role for both WASP-family members in promoting apoptotic F-actin-rich territory assembly and driving clustering-mediated caspase activation.

## DISCUSSION

Cytoskeletal studies related to apoptotic cell death have largely focused on changes in actin filament disassembly that drive terminal morphological phenotypes. However, roles for actin assembly proteins as active participants earlier in apoptosis have recently been uncovered, as the WASP-family members JMY and WHAMM both contribute to programmed cell death by enhancing cytosolic aspects of the intrinsic apoptosis pathway after p53-mediated cell cycle arrest (King et al., 2021). Our work now shows that JMY- and WHAMM-derived F-actin-rich territories promote apoptosis by creating a discrete intracellular environment for apoptosome biogenesis and caspase activation (Figure 7). These findings reveal new functions for the actin cytoskeleton in controlling the spatiotemporal progression of apoptosis, and create a framework for understanding how dynamic cytoskeletal responses organize specific signaling pathways.

**Figure 7.**
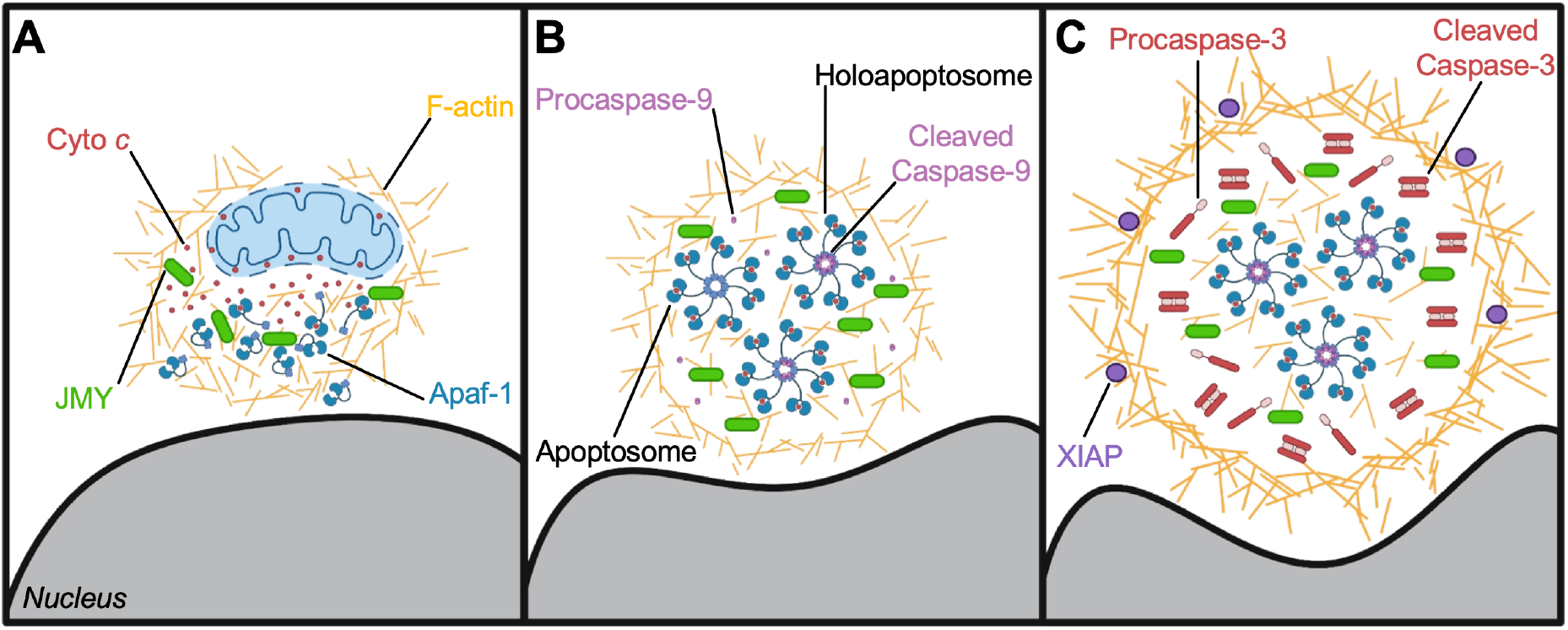
Model for apoptotic F-actin-rich territory assembly in coordinating apoptosome biogenesis and caspase activation. **(A)** Mitochondrial outer membrane permeabilization and release of cyto c (red circles) into the cytosol result in recruitment of JMY (green) and Arp2/3 complex-mediated assembly of actin filaments (gold) in a juxtanuclear region. WHAMM (not depicted) enhances actin polymerization and branching, and the early F-actin-rich territory sequesters cyto c and enables interactions with Apaf-1 (blue monomers). **(B**) F-actin remodeling and territory maturation creates a microenvironment conducive to the biogenesis and concentration of apoptosomes (blue/red heptamers). Initiator Procaspase-9 (pink) incorporates into holoapoptosomes (heptamers with purple hubs), which are retained within the territory. **(C)** The F-actin-rich territory increases in density and acts as a subcellular compartment that optimizes holoapoptosome-mediated processing of executioner Procaspase-3 (red monomers) into active Cleaved Caspase-3 (red dimers). F-actin functions to cluster the caspase activation process in the internal portion of the territory, while dense peripheral networks restrict accessibility of the caspase inhibitor XIAP (purple ovals). Eventually, active Caspase-3 achieves high enough quantities to bypass XIAP, escape the territory, and trigger the rapid proteolysis of substrates throughout the rest of the cell (not shown).

JMY has the ability to shuttle between the nucleus and cytosol (Zuchero et al., 2012), and we found that endogenous JMY is primarily nuclear both at steady state and after etoposide-induced DNA damage. Nevertheless, the cytosolic abundance of JMY increased significantly under the latter condition, a phenotype that is due to the formation of a juxtanuclear cluster of JMY puncta. JMY has previously been observed to form cytosolic puncta that associate with F-actin, and here we show that the accumulation of JMY puncta is promptly followed by the assembly and reorganization of F-actin into a unique cytoskeletal territory.

WHAMM and the Arp2/3 complex also localize throughout the JMY-containing F-actin territories, revealing that these structures contain the specific members of the actin assembly machinery that were previously found to be important for intrinsic apoptosis (King et al., 2021). In contrast, STRAP, a known inhibitor of JMY-mediated actin assembly (Hu and Mullins, 2019), and microtubules, which can repress WHAMM-mediated Arp2/3 activation (Shen et al., 2012), are absent from the F-actin-rich regions. These data are consistent with the idea that JMY and WHAMM stimulate Arp2/3-mediated actin assembly to construct pro-apoptotic territories that behave like compartments for segregating distinct subcellular components.

A critical point in the cellular commitment to apoptosis, mitochondrial permeabilization, is also somehow influenced by JMY and WHAMM. Mitochondrial outer membrane remodeling is caused by BCL-2 family proteins, which enable the export of cyto *c* to the cytosol. Some evidence indicates that this process initially occurs in a subset of mitochondria before propagating throughout the cell (Bhola et al., 2009; Rehm et al., 2009; Bock and Tait, 2020), although the signals that determine which mitochondria are permeabilized first and how a permeabilization wave may spread through the mitochondrial population are unclear. In etoposide-treated cells, JMY puncta were previously seen to partially colocalize with mitochondria-independent cytosolic cyto *c*, and JMY was found to be required for the formation of the cyto *c* puncta themselves (King et al., 2021). These observations, when combined with our current findings that the JMY and cyto *c* structures are confined to a single juxtanuclear F-actin-rich territory, imply that the initial site of mitochondrial permeabilization and the location of territory assembly may be one and the same (Figure 7A).

Apoptosomes, macromolecular scaffolds composed of cyto *c* and Apaf-1, are central players in intrinsic apoptosis and another focal point of regulation (Bratton and Salvesen, 2010; Yuan and Akey, 2013; Dorstyn et al., 2018). Biochemical and structural studies have defined the composition, configuration, and basic requirements for apoptosome biogenesis *in vitro*, but the mechanisms governing its assembly, organization, and activity are less well understood in cells. Cyto *c* and Apaf-1 are both enriched in F-actin territories, and JMY puncta also overlap with Apaf-1 puncta, albeit to a lesser extent than with cyto *c*. These data lead to the conclusion that apoptotic F-actin-rich territories compartmentalize the process of apoptosome biogenesis and that apoptosome formation in cells is dependent on JMY.

Apaf-1 monomers exist in a closed conformation, and cyto *c* binding triggers structural changes that allow the assembly of apoptosomes composed of 7 Apaf-1 molecules and 7 cyto *c* subunits arranged in a wheel-like structure with spokes that radiate from a central hub (Yuan et al., 2010; Yuan et al., 2013; Zhou et al., 2015; Cheng et al., 2016; Sahebazzamani et al., 2021). Apart from this structural information, the cellular mechanisms that dictate cyto *c* localization in the cytosol as well as its interactions with Apaf-1 are obscure. Our data begin to address this subject, as JMY and F-actin appear to mediate the early sequestration of mitochondria-released cyto *c* prior to its incorporation into apoptosomes. How Apaf-1 is recruited to territories is unknown, but the initial presence of cyto *c* within these nascent compartments likely facilitates Apaf-1 retention and integration into mature apoptosomes. Since most apoptotic F-actin-rich territories contain stoichiometric numbers of JMY, cyto *c*, and Apaf-1 puncta, we believe that the majority of cyto *c* and Apaf-1 structures visualized by immunofluorescence microscopy comprise groups of mature apoptosomes (Figure 7B).

Apoptosomes are important for the initiation of the caspase cleavage cascade by serving as platforms that interact with procaspase-9 monomers, resulting in their multimerization and activation (Li et al., 2017). Some evidence suggests that, after activation, caspase-9 remains bound to the apoptosome where it is able to process executioner caspases (Rodriquez and Lazebnik, 1999; Yin et al., 2006; Malladi et al., 2009; Yuan et al., 2011; Hu et al., 2014). However, the intracellular mechanisms underlying caspase recruitment to and activation by apoptosomes are not well characterized. Dense clusters of initiator caspase-9 and executioner caspase-3 are found in close proximity to collections of punctate JMY and apoptosomal proteins within F-actin-rich territories, leading us to conclude that these cytoskeletal structures are particularly conducive to apoptosome-mediated caspase activation.

The high frequencies with which F-actin territories surrounded JMY puncta alone or in conjunction with apoptosomal components likely places JMY and territory formation early in this signaling process. Nevertheless, the localization of initiator and active executioner caspases within F-actin territories shows that these apoptosome-enriched areas are maintained through the stage of the intrinsic pathway in which the caspase cleavage cascade is initiated. Aligning with a compartmentalization role for the territory, the caspase-inhibitor XIAP was restricted from accessing the internal portion where the majority of JMY, cyto *c*, and Apaf-1 were located. Thus, the F-actin-rich territories serve to concentrate both apoptosomes and caspases while shielding the caspases from a prominent inhibitor. As a consequence, such cytoskeletal rearrangements create a microenvironment that ensures a rapid and efficient transition from the biogenesis of apoptosomes to the induction of the caspase cascade (Figure 7C). We hypothesize that the temporary physical protection provided to the active caspases in the center of the territory permits them to reach quantities that can eventually overwhelm the peripheral inhibitors, escape the territory, and execute a sudden and complete demolition of the cell.

JMY requires its actin- and Arp2/3-binding domains to enable caspase-3 cleavage (King et al., 2021), and our current work demonstrates that JMY-mediated assembly of F-actin territories results in not only a more frequent triggering of executioner caspase activation on a per-cell basis, but that denser actin networks within the cell drive more potent processing of caspase-3 to its cleaved active form. The use of single and double knockout cells indicates that JMY and WHAMM each contribute to the amount of F-actin comprising territories. Based on the severity of mutant cell phenotypes, JMY is pivotal for creating these cytoskeletal structures, whereas WHAMM may increase the density of actin filaments within them. In addition to producing greater amounts of cleaved executioner caspases, F-actin-rich territories cause a distinct clustering of caspases, as subdued or absent F-actin territories in JMY- and WHAMM-deficient cells correspond to low and/or diffuse cleaved caspase signals. JMY and WHAMM therefore both function in organizing and clustering multiple apoptotic components within a single cytoskeletal compartment.

The proper spatiotemporal regulation of intracellular processes is achieved by compartmentalization in membrane-bound organelles or less well characterized membraneless organelles (Gabaldón and Pittis, 2015; Zhao and Zhang, 2020). Such subcellular partitioning enhances the rate and specificity of biochemical reactions by selectively increasing the proximity of individual proteins to positive regulators, while insulating these factors from negative regulators and competing reactions. The rapid creation of compartments *de novo* is well illustrated by the formation of viral assembly factories in infected cells (Miller and Krijnse-Locker, 2008; Schmid et al., 2014). Moreover, the biogenesis of membraneless organelles is often driven by multivalent interactions (Li et al., 2012; Gomes and Shorter, 2019), and the cytoskeleton plays important roles in their establishment, remodeling, and disassembly (Kaizuka et al., 2007; Banani et al., 2017; Koppers et al., 2020). Cytoskeletal filaments can also assemble into structures that segregate organelles and intracellular cargo, for example the actin cages that surround damaged mitochondria (Moore et al., 2016; Kruppa et al., 2018) or the actin cocoons that encapsulate bacteria-containing vacuoles (Kühn et al., 2020).

Consistent with the above themes, key members of the actin polymerization machinery and multiple pro-apoptotic proteins are selectively partitioned into F-actin-rich territories, which protect the actin nucleation factors from negative regulators and the caspases from competitive inhibitors. Determining how the apoptosomal and cytoskeletal components physically engage one another will shed light on the dynamic events that take place early in apoptosis, while investigations into the mechanisms by which caspases disseminate from the F-actin-rich territories will clarify how these proteases gain access to their diverse targets later in apoptosis. Thus, continuing to characterize the biogenesis, remodeling, dissolution, and broader impact of F-actin-rich territories could provide new avenues for understanding and modulating the cellular responses to genotoxic or other pro-apoptotic stressors.

## MATERIALS AND METHODS

### Ethics statement

Research conducted with biological materials was approved by the UConn Institutional Biosafety Committee. This study did not include human subjects or live animals.

### Cell culture

Cell lines are listed in Table S1, plasmids in Table S2, and key reagents in Table S3. U2OS cells (UC Berkeley cell culture facility) were cultured in Dulbecco’s Modified Eagle Medium (DMEM; Invitrogen) supplemented with Glutamax, 10% fetal bovine serum (FBS) and antibiotic-antimycotic. HAP1 cell derivatives (Horizon Genomics) were described previously (King et al., 2021), and were cultured in Iscove’s Modified Dulbecco’s Medium (IMDM; Invitrogen) supplemented with 10% FBS and penicillin-streptomycin. All cell lines were grown at 37°C in 5% CO_2_, and assays were performed using cells that had been in active culture for 2-10 trypsinized passages. Cells were treated with 5-10µM etoposide (Sigma Aldrich) diluted from an initial stock of 50µM dissolved in media. Equivalent volumes of media without etoposide were used as controls.

### Transgene expression

Plasmids encoding GFP-tagged mouse JMY (King et al., 2021), LAP (localization and affinity purification) tagged human WHAMM (Campellone et al., 2008), and mCherry-tagged Lifeact (Ohkawa and Welch, 2018) were described previously. For stable expression of GFP-JMY derivatives, U2OS or JMY^KO-1A^ cells cultured in 12-well plates were transfected with 2-5µg of linearized plasmid encoding GFP or GFP-JMY derivatives using LipofectamineLTX (Invitrogen). 24h later, the cells were transferred to media containing 0.6 or 1.5mg/mL G418 for 12-16 days. Surviving colonies were collected, expanded in 24-well plates, 6-well plates, and ultimately T-25 flasks prior to cryopreservation. U2OS cells were further enriched for GFP^+^ cells using a FACSAria flow cytometer (BD Biosciences). Upon re-animation, G418 concentrations were reduced to 350µg/mL in 6-well plates. The cell populations were subjected to experimental manipulations within 5 passages in media containing 350µg/mL G418. For transient expression of Lifeact-mCherry or LAP-WHAMM, U2OS cells were transfected in 6-well plates with 500ng of DNA 24h prior to reseeding onto 12mm glass coverslips in 24-well plates or into 35mm glass-bottom microwell dishes.

### Immunoblotting

To prepare whole cell extracts, cells were washed with phosphate-buffered saline (PBS), collected, and pelleted via centrifugation. Cell pellets were resuspended in lysis buffer (20mM HEPES pH 7.4, 100mM NaCl, 1% Triton X-100, 1mM Na_3_VO_4_, 1mM NaF, plus 1mM PMSF, and 10µg/ml each of aprotinin, leupeptin, pepstatin, and chymostatin), diluted in SDS-PAGE sample buffer, boiled, centrifuged, and subjected to SDS-PAGE before transfer to nitrocellulose membranes (GE Healthcare). Membranes were blocked in PBS + 5% milk (PBS-M) before being probed with primary antibodies (Table S3) diluted in PBS-M overnight at 4°C plus an additional 2-3h at room temperature. Membranes were rinsed twice with PBS and washed thrice with PBS + 0.5% Tween-20 (PBS-T). Membranes were then probed with secondary antibodies conjugated to IRDye-800, IRDye-680, or horseradish peroxidase (Table S3) and diluted in PBS-M. Membranes were again rinsed with PBS and washed with PBS-T. Blots were imaged using a LI-COR Odyssey Fc imaging system. Band intensities were determined using the Analysis tool in LI-COR Image Studio software, and quantities of proteins-of-interest were normalized to tubulin, actin, and/or GAPDH loading controls.

### Immunostaining

For immunofluorescence microscopy, approximately 1.5-2.5x10^5^ cells were seeded onto 12mm glass coverslips in 24-well plates and allowed to grow for 24h. After control or etoposide treatments, cells were washed with PBS, fixed in 2.5% or 3.7% paraformaldehyde (PFA) in PBS for 30min, washed, permeabilized with 0.1% TritonX-100 in PBS, washed, and incubated in blocking buffer (1% FBS + 1% bovine serum albumin (BSA) + 0.02% NaN_3_ in PBS) for a minimum of 15min. Cells were probed with primary antibodies (Table S3) diluted in blocking buffer for 45min. Cells were washed and treated with AlexaFluor-conjugated secondary antibodies, DAPI, and/or AlexaFluor-conjugated phalloidin (Table S3) for 45min, followed by washes and mounting in Prolong Gold anti-fade reagent (Invitrogen). For live imaging, approximately 10^6^ cells were seeded into 35mm glass-bottom dishes (MatTek) and allowed to grow for 24h prior to etoposide treatments.

### Fluorescence microscopy

All fixed and live images were captured using a Nikon Eclipse T*i* inverted microscope with Plan Apo 100X/1.45, Plan Apo 60X/1.40, or Plan Fluor 20x/0.5 numerical aperture objectives using an Andor Clara-E camera and a computer running NIS Elements software. Fixed cells were viewed in multiple focal planes, and Z-series were captured at 0.2-0.4µm steps. Images presented in the figures represent one slice or two-slice projections. Live cell imaging was performed in a 37°C chamber (Okolab) using the 100X objective, and images were captured at 2min intervals. All images were processed and analyzed using ImageJ/FIJI software (Schindelin et al., 2012).

### Image processing and quantification

The ImageJ Cell Counter plugin was used to quantify the percentage of cells with JMY puncta, cleaved caspase-3 positivity, or F-actin-rich territories by manually counting the total number of cells in the DAPI channel and the number of cells that were positive for cytosolic punctate JMY staining in the JMY or GFP-JMY channel, the number of cells that were positive for cleaved caspase-3 staining, or the number of cells that contained F-actin-rich territories. Similar quantifications were performed to determine the percentage of F-actin-rich territories that associated with JMY, GFP-JMY, cyto *c*, Apaf-1, or cleaved caspase-3 staining. The Cell Counter plugin was also used to quantify the percentage of cleaved caspase-3-positive cells that possessed F-actin-rich territories or exhibited clustered versus diffuse staining. Similar methods were used to perform JMY puncta overlap assays by identifying F-actin-rich territories in the phalloidin channel and manually counting the number of JMY puncta that did and did not overlap with cyto *c* or Apaf-1 puncta.

For analyses of total cellular JMY, cleaved caspase-3, or F-actin levels, the Selection tool was used in the phalloidin channel to select cells, and the Measure tool was used in the JMY, cleaved caspase-3, or phalloidin channel to measure the integrated density and area per cell, and then the integrated density was normalized to the cell area to generate the mean whole cell fluorescence. For analyses of nuclear JMY levels, the Threshold, Watershed, and Analyze tools were used in the DAPI channel to separate individual nuclei, the ROI Manager Measure tool was used in the JMY channel to measure the integrated density and area per nucleus, and then the integrated density was normalized to the nuclear area to generate the mean nuclear fluorescence. For analyses of cytoplasmic JMY levels, the nuclear integrated density or area was subtracted from the whole cell integrated density or area for individual cells, and then the cytosolic integrated density was normalized to the cytosolic area to generate the mean cytosolic fluorescence. The mean cytosolic fluorescence and mean nuclear fluorescence values were further normalized by dividing by their sum and multiplying by the mean whole cell fluorescence to derive the JMY intensities for each compartment on a per cell basis. For analysis of GFP-JMY or Lifeact-mCherry levels over time, the Selection tool was used in the mCherry and DAPI channel to select cytosolic or nuclear areas, and the Measure tool was used in the GFP or mCherry channel to measure the integrated density and area per selection. Then the integrated density was normalized to the selection area to generate the mean fluorescence for each timepoint and the average mean fluorescence for each channel was set to one.

Pixel intensity plots were generated using the Plot Profile tool. Lines were drawn through F-actin- or JMY-rich regions after identification in the phalloidin, cleaved caspase-3, or JMY channels, and the Plot Profile tool was used to measure the distance and integrated densities for each channel. The mean integrated density for each channel was set to one. For analyses of JMY, XIAP, and F-actin levels within F-actin-rich territories, the Selection tool was used in the phalloidin channel to set the territory area, and the Measure tool was used in the JMY, XIAP, and phalloidin channels to measure the integrated density per selection. Analyses of interior JMY, XIAP, or F-actin levels utilized the Enlarge Selection tool to shrink the Selected area by 2µm, and the Measure tool was used in the JMY, XIAP, and phalloidin channels to measure the integrated density. In measurements of JMY, XIAP, or F-actin levels at the territory periphery, the interior territory integrated density was subtracted from the whole territory integrated density for individual cells. The mean whole territory integrated density for each channel was set to one.

### Reproducibility and statistics

All conclusions were based on observations made from at least 3 separate experiments, and quantifications were based on data from 3 representative experiments, except where noted in the Figure Legends. The sample size used for statistical tests was the number of times an experiment was performed, except where noted in the Legends. Statistical analyses were performed using GraphPad Prism software. Statistics on data sets with 3 or more conditions were performed using ANOVAs followed by Tukey’s post-hoc test. P-values for data sets comparing 2 conditions were determined using unpaired t-tests. P-values for data sets with +/-scoring used Fisher’s exact test. P-values <0.05 were considered statistically significant.

### Supplemental material

Supplemental material includes 3 Tables describing biological and chemical reagents, 5 Figures of supporting experimental data, and 1 Video that accompanies the time-lapse images in Figure 1.

## ACKNOWLEDGEMENTS

We thank Wu He at the UConn Flow Cytometry Facility for cell sorting and L.T. Bear for support with experimental design. We also thank Katrina Velle, Nathan Leclair, and Campellone Lab members for their comments on this paper. KGC was supported by National Institutes of Health grants R01-GM107441 and K02-AG050774. The funders had no role in study design, data collection and analysis, decision to publish, or preparation of the manuscript. The authors declare no competing financial interests.

## AUTHOR CONTRIBUTIONS

Conceptualization, Data curation, Formal analysis, Investigation, Methodology, Validation, Visualization, Writing – original draft, Writing – review & editing: VLK and KGC. Funding acquisition, Project administration, Supervision: KGC.

## Notes

### Competing Interest Statement

The authors have declared no competing interest.

