## Supplement for "F-actin-rich territories coordinate apoptosome assembly and caspase activation during intrinsic apoptosis"

**Table S1. Cell Lines.**

| | | | | |
|---|---|---|---|---|
| ***Parental Cells*** | | | | |
| Cell Line | | Source | | |
| eHAP | | Horizon Genomics (C669) | | |
| HAP1 | | Horizon Genomics (C631) | | |
| U2OS | | UC Berkeley Cell Culture Facility | | |
| ***HAP1 Derivatives*** | | | | |
| KO Cell Line | | Mutation | Predicted AAs | Source |
| JMY$^{KO-1A}$ | | 17bp deletion in exon 1 of 11 | 145/988, 4 post-shift | Horizon Genomics (HAP1_JMY_28380-03) King et al., 2021 |
| JMY$^{KO-2}$ | | 2bp deletion in exon 2 of 11 | 362/988, 3 post-shift | Horizon Genomics (HZGHC002631c007) King et al., 2021 |
| ***eHAP Derivatives*** | | | | |
| KO Cell Line | | Mutation | Predicted AAs | Source |
| WHAMM$^{KO-2}$ | | 10bp deletion in exon 2 of 10 | 204/809, 7 post-shift | Horizon Genomics Mathiowetz et al., 2017 King et al., 2021 |
| WHAMM$^{KO-4}$ | | 7bp deletion in exon 4 of 10 | 321/809, 17 post-shift | Horizon Genomics (HZGHC001060c001) King et al., 2021 |
| WHAMM/JMY$^{DKO-1}$ | WHAMM: | 10bp deletion in exon 2 of 10 | 204/809, 7 post-shift | Horizon Genomics (HZGHC004884C012) King et al., 2021 |
| | JMY: | 16bp deletion in exon 1 of 11 | 112/998, 64 post-shift | |
| WHAMM/JMY$^{DKO-2}$ | WHAMM: | 10bp deletion in exon 2 of 10 | 204/809, 7 post-shift | Horizon Genomics (HZGHC004884C007) King et al., 2021 |
| | JMY: | 35bp deletion in exon 2 of 11 | 362/998, 33 post-shift | |

**Table S2. Plasmids.**

| ***Plasmids*** | | | | | |
|---|---|---|---|---|---|
| Description | Vector | Species | AA | R.E. Sites | Source |
| pGFP (pKC425) (vector) | pCDNA3::GFP | N/A | N/A | N/A | Campellone et al., 2008 |
| pGFP-JMY | pCDNA3::GFP | Mouse | 1-983 | BamHI-NotI | King et al., 2021 |
| pGFP-JMY(ΔCA) | pCDNA3::GFP | Mouse | 1-941 | BamHI-NotI | King et al., 2021 |
| pGFP-JMY(ΔWWW) | pCDNA3::GFP | Mouse | 1-856/ 934-983 | BamHI-NotI | King et al., 2021 |
| pLAP (pKC-LAP-C1) (vector) | pIC113::HisGFP-S | N/A | N/A | N/A | Campellone et al., 2008 |
| pLAP-WHAMM | pIC113::HisGFP-S | Human | 1-809 | KpnI-BamHI | Campellone et al., 2008 |
| pACT-Lifeact-MCH | pACT::MCH | N/A | N/A | SpeI-BglII | Ohkawa and Welch, 2018 |

**Table S3. Immunofluorescence and Immunoblotting Reagents.**

***Primary Antibodies (Immunofluorescence)***

| *Target* | *Probe* | | *Conc.* | *Identifier* |
|---|---|---|---|---|
| Apaf-1 (Fig 3, 4) | anti-Apaf-1 | Mouse | 1:250 | R&D Systems (MAB868) |
| Arp3 (Fig 2) | anti-ARP3 | Mouse | 1:1,000 | Sigma (A5979) |
| ArpC2 (Fig 2, 3) | anti-p34-Arc | Rabbit | 1:1,000 | EMD Millipore (07-227-I) |
| Caspase-9$^{\text{Pro \& Cleaved}}$ (Fig S4) | anti-Total Caspase-9 | Rabbit | 1:500 | Cell Signaling Technology (9502) |
| Caspase-3$^{\text{Pro \& Cleaved}}$ (Fig S4) | anti-Total Caspase-3 | Mouse | 1:500 | Cell Signaling Technology (9668) |
| Caspase-3$^{\text{Cleaved}}$ (Fig 4, 5, 6, S4) | anti-Cleaved Caspase-3 | Rabbit | 1:1,000 | Cell Signaling Technology (9664) |
| Cortactin (Fig S2) | anti-Cortactin | Mouse | 1:2,000 | EMD Millipore (05-180) |
| Cytochrome *c* (Fig 3, 4) | anti-Cytochrome *c* | Mouse | 1:500 | Cell Signaling Technology (12963) |
| Golgi (Fig S3) | anti-GM130 | Mouse | 1:1,000 | BD Transduction Labs (610822) |
| JMY (Fig 1, 2, 3, 4, S2, S3, S4) | anti-JMY | Rabbit | 1:1,000 | Proteintech (25098-1-AP) |
| Mitochondria (Fig S3) | anti-AIF | Rabbit | 1:1,000 | Cell Signaling Technology (5318) |
| N-WASP (Fig S2) | anti-N-WASP | Guinea Pig | 1:1,000 | Duleh et al., 2010 |
| p53 (Fig S3) | anti-p53 | Rabbit | 1:1,000 | Proteintech (10442-1-AP) |
| STRAP (Fig 2) | anti-STRAP | Mouse | 1:1,000 | Proteintech (66712-1-Ig) |
| Tubulin (Fig 2) | anti-Beta-Tubulin | Mouse | 1:2,000 | DSHB (E7) |
| WASH (Fig S2) | anti-WASH | Rabbit | 1:1,000 | Duleh et al., 2010 |
| WAVE2 (Fig S2) | anti-WAVE2 | Rabbit | 1:1,000 | Cell Signaling Technology (3659) |
| WHAMM (Fig 2, 3) | anti-WHAMM | Rabbit | 1:250 | Shen et al., 2012 |
| XIAP (Fig 4) | anti-XIAP | Mouse | 1:1,000 | Proteintech (66800-1-Ig) |

***Primary Antibodies (Immunoblotting)***

| *Target* | *Probe* | | *Conc.* | *Identifier* |
|---|---|---|---|---|
| Actin (Fig S1) | anti-Beta-Actin | Mouse | 1:10,000 | Proteintech (66009-1-Ig) |
| GAPDH (Fig S1) | anti-GAPDH | Mouse | 1:10,000 | Proteintech (60004-1-Ig) |
| GFP (Fig S1) | anti-GFP | Mouse | 1:1,000 | Santa Cruz (sc9996) |
| JMY (Fig S1) | anti-JMY | Rabbit | 1:1,000 | Proteintech (25098-1-AP) |
| Tubulin (Fig S1) | anti-Beta-Tubulin | Mouse | 1:10,000 | DSHB (E7) |

***Secondary Antibodies (Immunofluorescence)***

| *Target* | *Probe* | | *Conc.* | *Identifier* |
|---|---|---|---|---|
| Mouse IgG | Alexa 555 anti-mouse | Goat | 4 µg/ml | Life Technologies (e.g. A11029) |
| Rabbit IgG | Alexa 488, 555, 647 anti-rabbit | Goat | 4 µg/ml | Life Technologies (e.g. A11034) |
| Guinea Pig IgG | Alexa 555 anti-guinea pig | Goat | 4 µg/ml | Life Technologies (e.g. A11075) |

***Secondary Antibodies (Immunoblotting)***

| *Target* | *Probe* | | *Conc.* | *Identifier* |
|---|---|---|---|---|
| Mouse IgG | HRP anti-Mouse | Sheep | 1:10,000 | GE Healthcare (NXA931) |
| Rabbit IgG | HRP anti-Rabbit | Donkey | 1:10,000 | GE Healthcare (NA934V) |
| Mouse IgG | IRDye 680, 800 anti-Mouse | Donkey | 0.05 µg/ml | LI-COR (926-32212) |
| Rabbit IgG | IRDye 680, 800 anti-Rabbit | Donkey | 0.05 µg/ml | LI-COR (926-32213) |

| ***Molecular Probes (Fluorescence)*** | | | |
|---|---|---|---|
| *Target* | *Probe* | *Conc.* | *Identifier* |
| DNA (Fig 1, 5) | 4′,6-diamidino-2-phenylindole (DAPI) | 1 µg/ml | Invitrogen (D1306) |
| DNA (Fig 1) | Hoescht 33342 Solution | 2 µg/ml | Thermo Scientific (62249) |
| F-actin (Fig 2, 3, 4, 5, S2, S3, S4) | Alexa647-Phalloidin | 0.4 U/ml | Invitrogen (A22287) |
| F-actin (Fig 1, 6) | Alexa555-Phalloidin | 0.4 U/ml | Invitrogen (A34055) |
| F-actin (Fig 3, 4) | Alexa350-Phalloidin | 0.6 U/ml | Life Technologies (A22281) |

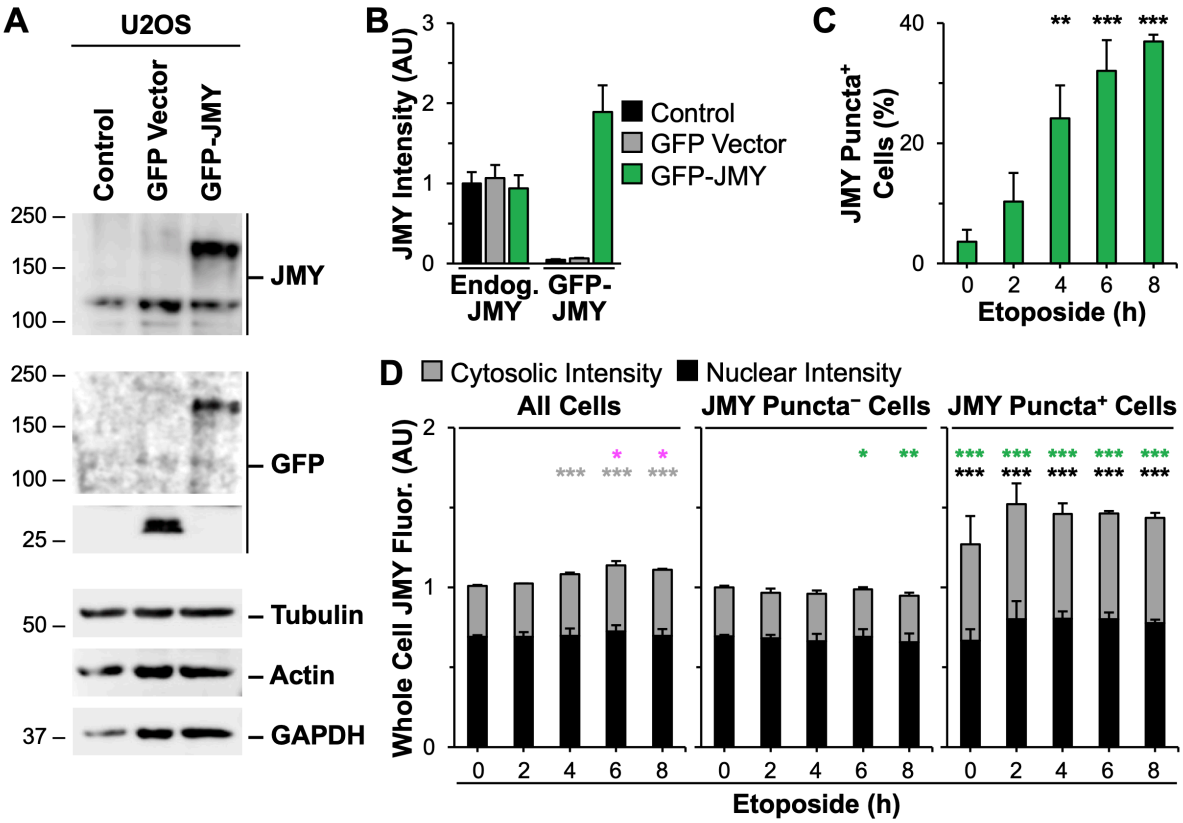


**Figure S1. JMY forms juxtanuclear clusters of puncta that are responsible for an increase in cytosolic JMY expression following DNA damage. (A-B)** U2OS cell lysates from control untransfected cells or cells stably expressing a GFP vector or GFP-JMY were immunoblotted with antibodies to JMY, GFP, tubulin, actin, and GAPDH. For quantification, the JMY band intensities for endogenous JMY and GFP-JMY were normalized to the loading control intensities and the endogenous control value for JMY was set to 1. Each bar is the mean intensity ±SD from 3 blots. AU = Arbitrary Units. **(C-D)** Cells were treated as in Figure 1A-C. In (C), data from Figure 1A were replotted so each bar is the mean±SD from 3 experiments (n = 310-348 cells per point) and black significance stars refer to comparisons to the 0h timepoint. In (D), data from Figure 1C were replotted to show the mean whole cell JMY fluorescence in the full bar. The gray portion of the bar is the mean cytosolic JMY intensity, and the black portion is the mean nuclear JMY intensity. Each bar is the mean±SD from 3 experiments (n = 310-348 cells per timepoint). Magenta significance stars refer to comparisons of the total population whole cell fluorescence values at each timepoint to the 0min timepoint, and gray significance stars are to the cytosolic intensity value. Green significance stars refer to comparisons of whole cell fluorescence values for puncta-negative or puncta-positive populations to the total population at each timepoint and black significance starts are to the cytosolic intensity values. *p<0.05; **p<0.01; ***p<0.001 (ANOVA, Tukey post-hoc tests).

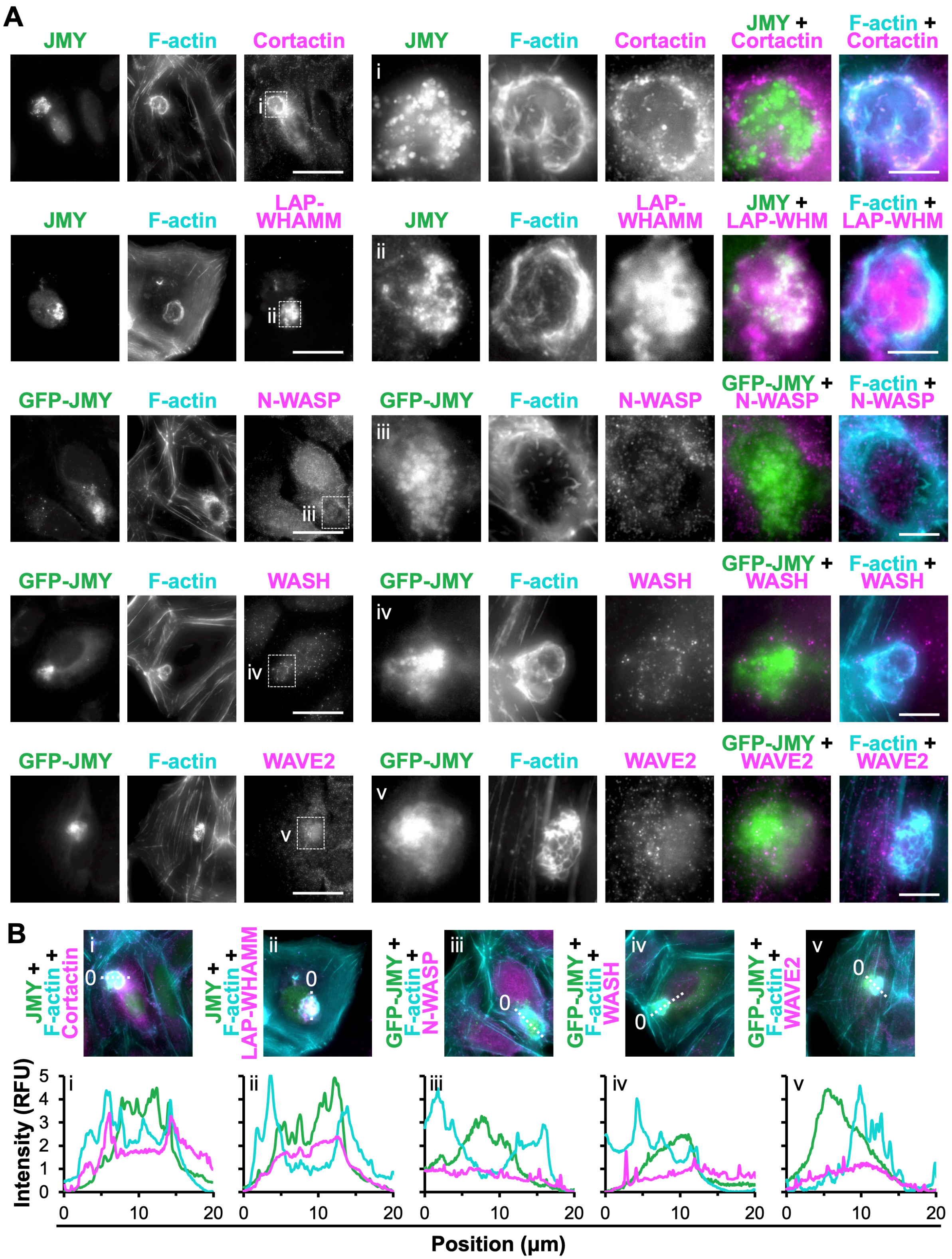


**Figure S2. WHAMM, but not representatives of other subgroups within the WASP family, is enriched within apoptotic F-actin-rich territories. (A)** U2OS cells or U2OS cells transfected with plasmids encoding GFP-JMY (green) or LAP-WHAMM (magenta) were treated *(continued)* with 10µM etoposide for 6h, fixed, and stained with phalloidin (F-actin; cyan) and antibodies to JMY (green), Cortactin, N-WASP, WASH, or WAVE2 (each in magenta). Magnifications (i-ii) depict F-actin territories with JMY, Cortactin, and WHAMM, while (iii-v) depict F-actin territories without enrichment of N-WASP, WASH, or WAVE2. Scale bars, 25µm, 5µm. **(B)** 20µm lines were drawn through the images in (A) using ImageJ to measure the pixel intensity profiles. RFU = relative fluorescence units.

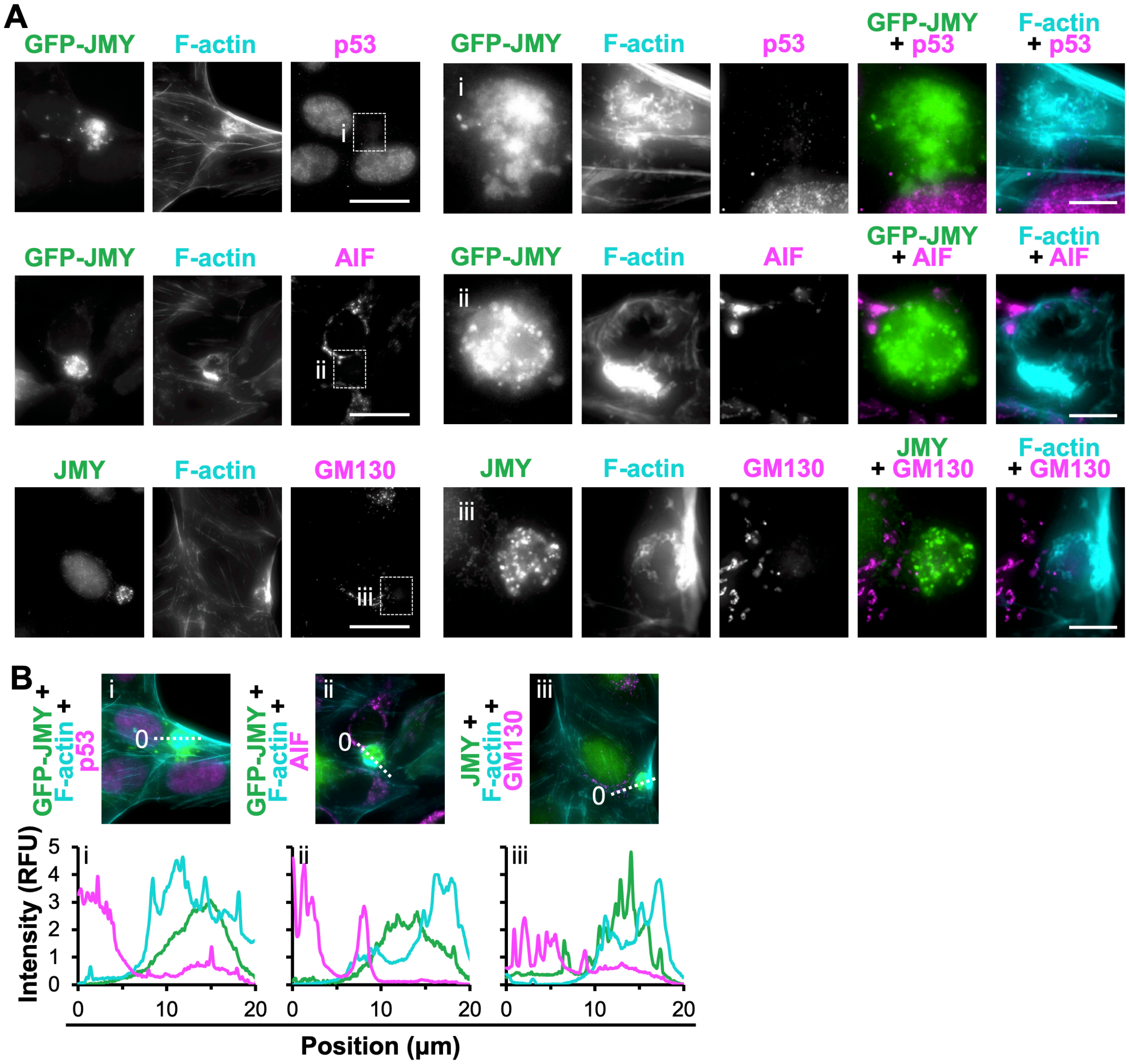


**Figure S3. p53, mitochondria, and the Golgi are not recruited to apoptotic F-actin-rich territories. (A)** U2OS cells or U2OS cells expressing GFP-JMY (green) were treated with etoposide for 6h, fixed, and stained with phalloidin (F-actin; cyan) and antibodies to JMY (green), p53, AIF, or GM130 (each in magenta). Magnifications (i-iii) depict JMY and F-actin territories lacking p53, mitochondria (AIF), or Golgi (GM130). Scale bars, 25µm, 5µm. **(B)** 20µm lines were drawn through the images in (A) using ImageJ to measure the pixel intensity profiles.

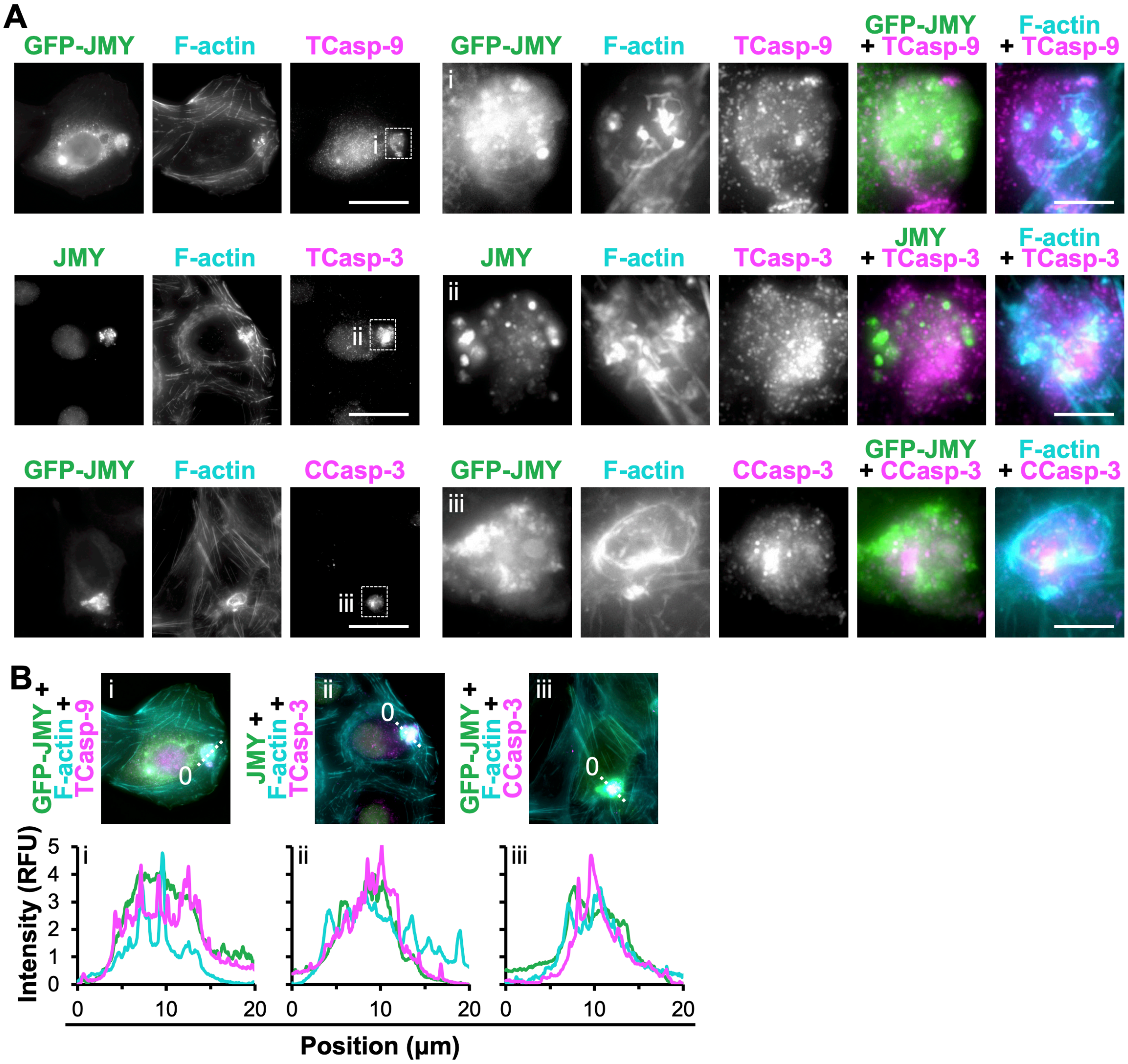


**Figure S4. Initiator caspase-9 and executioner caspase-3 are found within apoptotic F-actin-rich territories. (A)** U2OS cells or U2OS cells expressing GFP-JMY (green) were treated with etoposide for 6h, fixed, and stained with phalloidin (F-actin; cyan), antibodies to JMY (green), and either antibodies that recognize total caspase-9 including uncleaved and the large fragment that results from cleavage at Asp315 (TCasp-9; magenta), antibodies that recognize total caspase-3 including uncleaved and the large fragment resulting from cleavage at Asp175 (TCasp-3; magenta), or antibodies that recognize only active caspase-3 cleaved at Asp175 (CCasp-3; magenta). Magnifications (i-iii) depict JMY and F-actin territories containing total caspase-9, total caspase-3, and cleaved caspase-3. Scale bars, 25μm, 5μm. **(B)** 20μm lines were drawn through the images in (A) using ImageJ to measure the pixel intensity profiles.

A

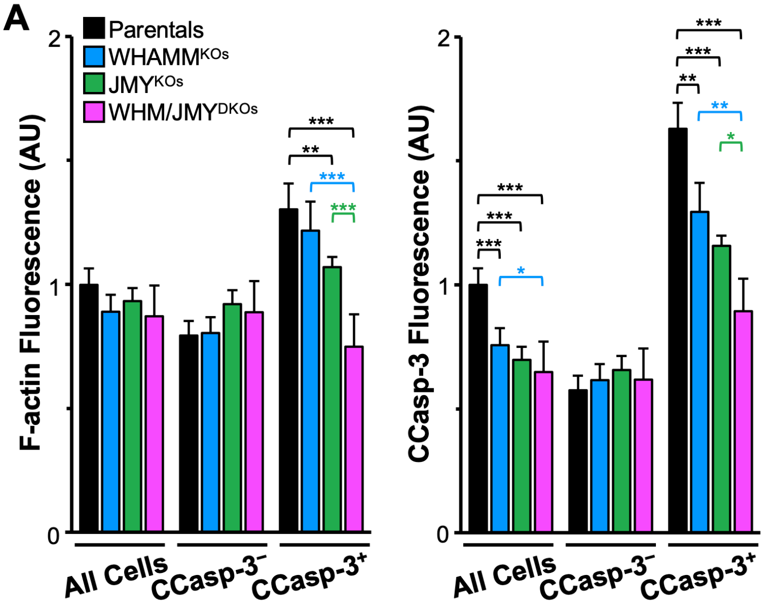


**Figure S5. The amount of F-actin and caspases within apoptotic cells is enhanced by WHAMM. (A)** Cells were treated as in Figure 6A. Whole cell fluorescence values for F-actin and cleaved caspase-3 were measured in individual cells using ImageJ and normalized to the parental samples. Values for the CCasp-3-postitve cell population alone appears in Figure 6G. Each bar is the mean±SD from 3 experiments (n = 607-866 cells per sample). $*p<0.05$; $**p<0.01$; $***p<0.001$ (ANOVA, Tukey post-hoc tests).

**Video S1. Time-lapse movie showing the formation of punctate GFP-JMY structures and an F-actin-rich territory.** Live U2OS cell images of GFP-JMY (green) and Lifeact-mCherry (magenta) fluorescence were captured at 2min intervals 180-260min after exposure to 10μM etoposide (related to Figure 1). Playback rate = 10 frames per second. Scale bar, 20μm.
